# Generalising uncertainty improves accuracy and safety of deep learning analytics applied to oncology

**DOI:** 10.1101/2022.07.14.500142

**Authors:** Samual MacDonald, Helena Foley, Melvyn Yap, Rebecca L. Johnston, Kaiah Steven, Lambros T. Koufariotis, Somwya Sharma, Scott Wood, Venkateswar Addala, John V. Pearson, Fred Roosta, Nicola Waddell, Olga Kondrashova, Maciej Trzaskowski

## Abstract

Trust and transparency are critical for deploying deep learning (DL) models into the clinic. DL application poses generalisation obstacles since training/development datasets often have different data distributions to clinical/production datasets that can lead to incorrect predictions with underestimated uncertainty. To investigate this pitfall, we benchmarked one pointwise and three approximate Bayesian DL models used to predict cancer of unknown primary with three independent RNA-seq datasets covering 10,968 samples across 57 primary cancer types. Our results highlight simple and scalable Bayesian DL significantly improves the generalisation of uncertainty estimation (e.g., p-value = 0.0013 for calibration). Moreover, we demonstrate Bayesian DL substantially improves accuracy under data distributional shifts when utilising ‘uncertainty thresholding’ by designing a prototypical metric that evaluates the expected (accuracy) loss when deploying models from development to production, which we call the Area between Development and Production curve (ADP). In summary, Bayesian DL is a hopeful avenue of research for generalising uncertainty, which improves performance, transparency, and therefore safety of DL models for deployment in real-world.

---

Recent advances in deep learning (DL) have led to the rapid development of diagnostic and treatment support applications in various aspects of healthcare, including oncology [1]–[4]. The proposed applications of DL utilise a range of data modalities, including MRI scans [5], CT scans [6], histopathology slides [7], genomics [8], transcriptomics [9], [10], and most recently, integrated approaches with various data types [11], [12]. In general, studies using DL show excellent predictive performance, providing hope for successful translation into clinical practice [13], [14]. However, prediction accuracy in DL comes with potential pitfalls which need to be overcome before wider adoption can be eventuated [15].

The lack of transparency over prediction reliability is one challenge for implementing DL [16]. One approach to overcome this is by providing uncertainty estimates about a model’s prediction [17, p. 20], [18], enabling better-informed decision making. Another obstacle relates to the assumptions made about data when transitioning from training to real-world applications. In standard DL practice, during the ‘development’ stage, models are trained and validated on data prepared to satisfy the assumption of independent and identically distributed (IID) data, meaning that model would be applied to make predictions on the data that are independently drawn and come from the same distribution as the training data. However, this assumption is frequently violated when models are deployed in ‘production’ (i.e. real-world application), when confounding variables cause distributional shifts that push data out-of-distribution (OOD) [19]. For oncology applications, such confounding variables can include technical differences in how the data are collected (e.g. batch effects, differences in sequencing depth or library choice for genomic and transcriptomic data; differences in instrumentation and imaging settings for medical imaging data), as well as biological differences (e.g. differences in patient demographics or a data class unseen during model development). The consequences from OOD data include inaccurate predictions coupled with underestimated uncertainties, which together result in the model’s overconfidence from distributional shifts, or what we call ‘shift-induced’ overconfidence [20]–[22]. Consequently, implementation of DL into clinical practice (i.e., production) requires that models are robust (i.e., generalise) to distributional shifts and provide correct predictions with calibrated uncertainties.

Methods to address DL overconfidence in production exist, albeit with different limitations. Repeated retraining of deployed models on new production data is beneficial for accuracy, but introduces new risks such as over-computation or catastrophic forgetting, whereby DL models lose performance on original training/development data [23], [24]. Using tracking metrics such as accuracy can help inform ML engineers about the DL reliability, although such metrics are only available retrospectively. A key pitfall for these methods are that they are reactive and not proactive.

One proactive approach for managing risks from DL overconfidence in production is with ‘uncertainty thresholding’, whereby only predictions with uncertainties below a threshold are accepted (to increase accuracy). Importantly, a DL model’s uncertainty threshold is established with development (IID) data. Thus, when the model is deployed to (OOD) production data it becomes overconfident. Therefore, the uncertainty threshold (established in development) corresponds to higher error-rate in production, which is a problem if expectations (between healthcare professionals and engineers) are set during the development phase of a project’s life cycle. To address this problem, post-hoc methods exist that calibrate uncertainty (e.g., with ‘Temperature scaling’; [25]). However, while post-hoc calibration effectively controls overconfidence in IID data [25], it fails to do so (proactively) in OOD data [21], [22]. Despite the notable theoretical and empirical research towards generalising DL uncertainties from OOD data [26], [27], shift-induced overconfidence is yet to be sufficiently addressed in practice.

In this study, we aim to address the safety and performance concerns of shift-induced overconfidence (i.e., the generalisation of uncertainty). We establish theoretical and empirical evidence of the phenomenon using a case study that predicts cancer of origin with transcriptomic data. Cancer of origin prediction has been an active application area for DL [24], [28]–[30], since accurate diagnosis is critical for the treatment of cancers of unknown primary (CUP), i.e. metastatic cancers in which the primary cancer site cannot be reliably determined. We investigate this case study’s dataset, with simple and accessible (i.e. relevant) DL techniques that help generalise uncertainty. Finally, we establish a prototypical metric, ADP, alongside a small discussion about how it may be helpful in a clinical setting.

## Results

### Bayesian model benchmarking approach to predict cancer of unknown primary

The primary DL task was to predict the tissue of origin (primary cancer type) of cancer samples using transcriptomic data. We used transcriptomic data from TCGA of primary cancer samples corresponding to 32 primary cancer types as model ‘development’ data: training (n=8,202; [31]) and validation IID data (n=1,434; Supplementary Table 1). The test data were OOD (representing ‘production’), providing a platform for benchmarking resilience to overconfidence, and included TCGA metastatic samples (n=392; [32]), Met500 metastatic samples (n=479; [33]), and a combination of primary and metastatic samples from our own independent internal custom dataset, i.e. ICD (n=461; [34]–[42]; Fig. 1a, Supplementary Fig. 1). The distributional shifts in the test data were likely to be caused by several factors, including dataset batches, sample metastasis status (metastatic or primary) and whether the cancer type was absent during training (‘unseen’).

**Figure 1.**
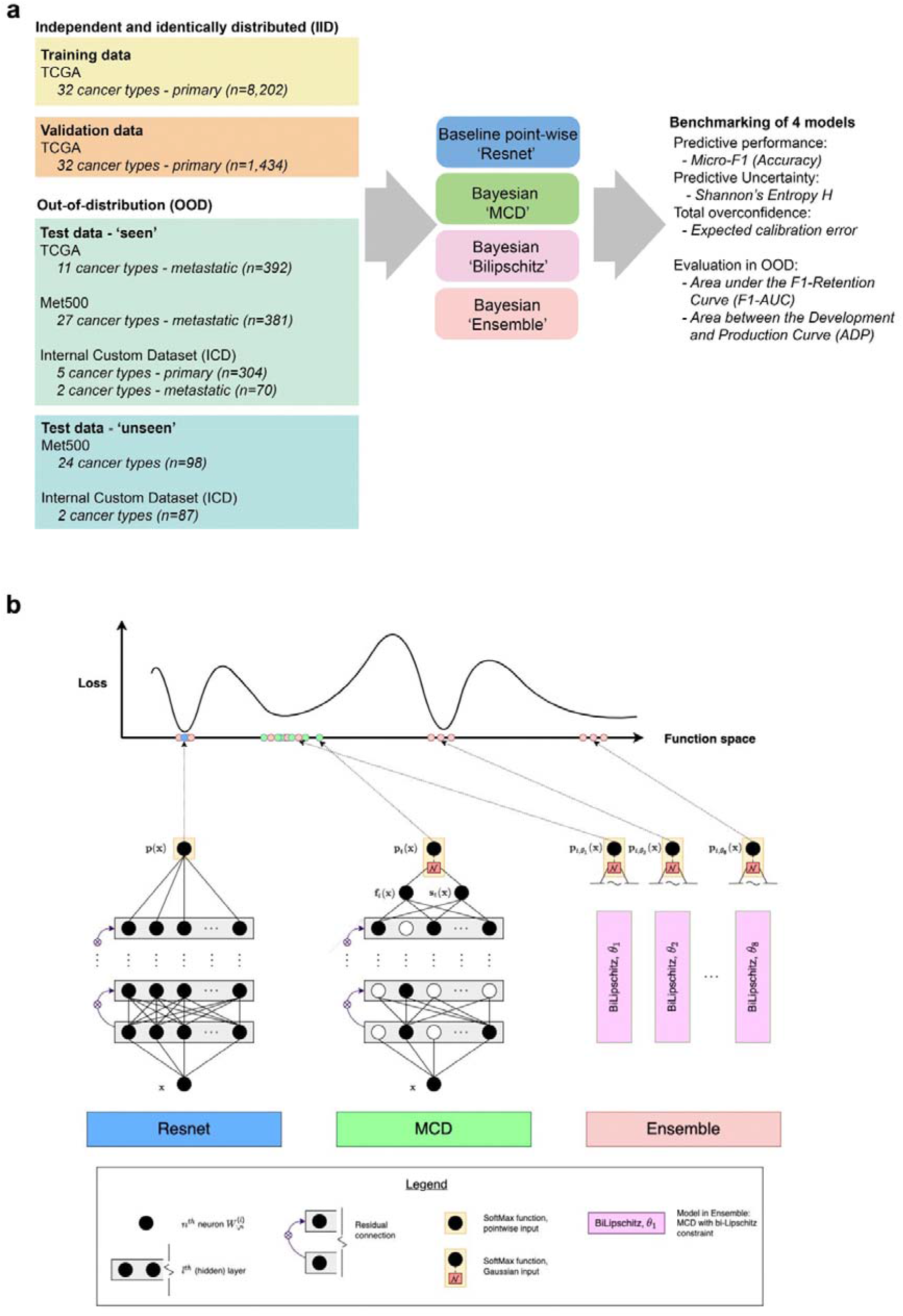
Overview of the study design. **a** Simplified study workflow. TCGA primary cancer types comprised the training and IID validation data. OOD test data comprised of the TCGA (metastatic cancer types), Met500 and ICD datasets, which included primary, metastatic and ‘unseen’ cancer types. **b** Schematic overview of the four tested models: pointwise Resnet (Resnet), Resnet extended with Monte Carlo Dropout (MCD), MCD extended with bi-Lipschitz constraint (Bilipschitz), and an ensemble of Bilipschitz models (Ensemble). Note, Resnet represents a single point in function space (blue dot), while two Bayesian models (MCD and Bilipschitz) represent a distribution within a single region in function space (green dots). The Ensemble represents a collection of distributions centred around different modes (red dots).

We aimed to evaluate if three simple ‘distribution-wise’ (e.g. Bayesian) DL models (with Resnet architecture) improve performance and reduce shift-induced overconfidence compared to a pointwise baseline model (with identical Resnet architecture). To achieve this, we performed controlled benchmarking of the models over IID and OOD data (Fig. 1b). The Bayesian models were Monte Carlo Dropout approximation (‘MCD’) [43], MCD with smoothness and sensitivity constraints (‘Bilipschitz’) [44], [45], and an ensemble of Bilipschitz models (‘Ensemble’) [45]. The ways in which models differed were canonical: MCD modified Resnet by keeping Dropout during prediction, Bilipschitz modified MCD with spectral normalisation, Ensemble modified Bilipschitz by combining multiple models.

### Approximate Bayesian inference reduces shift-induced overconfidence for ‘seen’ classes in a primary cancer site context

The predictive performance of each model to predict primary tissue was assessed using micro-F1 (equivalent to Accuracy; abbreviated F1). For the IID validation data, the difference between the highest and lowest ranking models was 0.28% (97.07% for Resnet and 96.79% for Ensemble, respectively; Fig. 2a, Supplementary Fig. 2-5). As expected, F1 scores dropped for the OOD test set across all four models, with a 1.74% difference between the highest and lowest ranking models (82.04% for Ensemble and 80.30% for Resnet, respectively; Fig. 2b, Supplementary Fig. 6-9). All models had higher predictive uncertainties (Shannon’s entropy ***H***) for OOD, relative to IID data (Fig. 2b). Uncertainties were significantly higher for all approximate Bayesian models (MCD, Bilipshitz, and Ensemble) relative to (pointwise) Resnet (*P* < 0.0001). Moreover, overconfidence in OOD data was evident for the Resnet and MCD models since their binned accuracies (i.e. the correct classification rates within bins delineated by the confidence scores) were consistently lower than corresponding confidence scores (Fig. 2c). The expected calibration errors (ECEs) for OOD data ranged between 5% for Ensemble and Bilipschitz and 16% for Resnet (Fig. 2c). Estimation of overconfidence as an absolute error was negligible across all models for IID data, with high amounts of overconfidence for OOD data, highlighting the shift-induced overconfidence when transitioning from IID to OOD data (Fig. 2d). Furthermore, Resnet had significantly higher overconfidence than MCD (p-value < 0.01), Bilipschitz (p-value < 0.001), and Ensemble (p-value < 0.001) for OOD data but not IID data. This shows that the shift-induced overconfidence in pointwise DL models can be reduced with simple (approximate) Bayesian inference.

**Figure 2.**
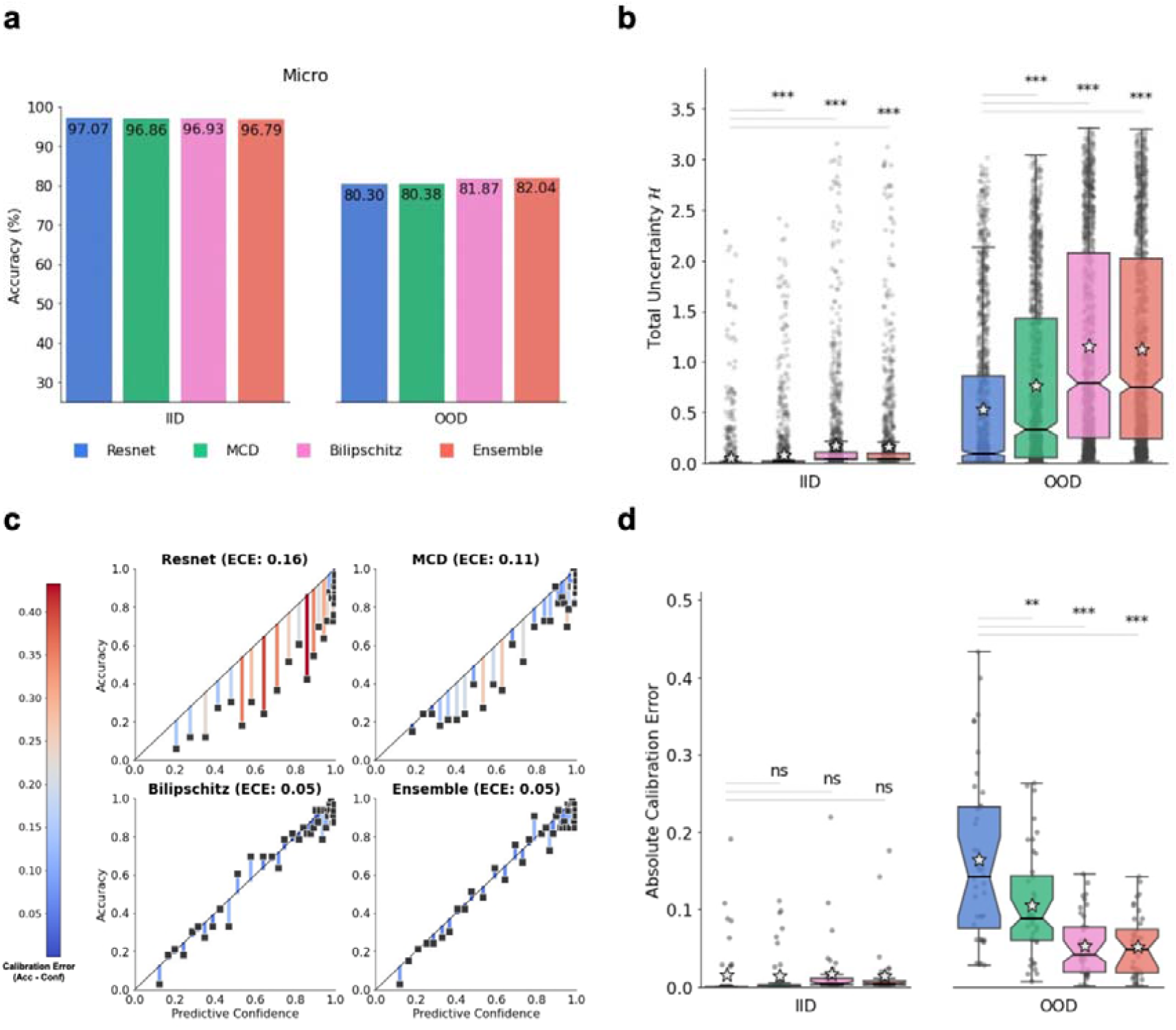
Out-of-distribution overconfidence of a pointwise baseline Resnet model and three simple Bayesian models on ‘seen’ data. **a** Micro-F1 score (i.e. Accuracy) of all models on the IID validation data (left) and on ‘seen’ OOD data (right). Accuracy for (IID) validation data was controlled with early stopping. **b** Box plot of each model’s predictive uncertainty (Shannon’s Entropy, *H*) for individual samples on IID data (left) and on ‘seen’ OOD data (right). Sample median is depicted by horizontal line, while the sample mean is depicted by the grey star. Statistical significance (single-sided Wilcoxon rank-sum) between baseline and each Bayesian model are marked with denoted *, **, ***, for p-value < 0.05, p-value < 0.01, and p-value < 0.001, respectively. **c** Each model’s confidence vs accuracy of each ECE-bin on ‘seen’ OOD data. The black diagonal lines illustrate perfect calibration, i.e., no overconfidence. ECE value for each model shown in parentheses. The residuals are colour-coded by the (left) colour scale and represent the difference between confidence and accuracy for each bin. **d** Box plot of each model’s absolute calibration error of individual samples on IID data (left) and ‘seen’ OOD data (right). Statistical significance (single-sided Wilcoxon rank-sum) between baseline and each Bayesian model are marked with denoted *, **, ***, for p-value < 0.05, p-value < 0.01, and p-value < 0.001, respectively.

### Prediction overconfidence for ‘unseen’ classes explained by related primary cancer types

Classes absent from training (‘unseen’) cannot have correct predictions, and prediction uncertainties should be higher compared to ‘seen’ classes. As expected, mean total uncertainties were higher for ‘unseen’ classes for all models (Fig. 3a). Moreover, approximate Bayesian models were significantly more uncertain with ‘unseen’ classes compared to Resnet (p-value < 0.01; Fig 3a). However, exceptions occurred across all models, where total uncertainty values were low, at both, class level, where predictions for a whole ‘unseen’ class consistently had low uncertainty, and sample level, where predictions for only some samples from a class had low uncertainty (Fig. 3b). We wanted to investigate whether any of the exceptions could be examples of ‘silent catastrophic failure’ (Supplementary Information – S4.2), a phenomenon where data are far from the training data’s support, resulting in incorrect yet extremely confident predictions [44]–[46].

**Figure 3.**
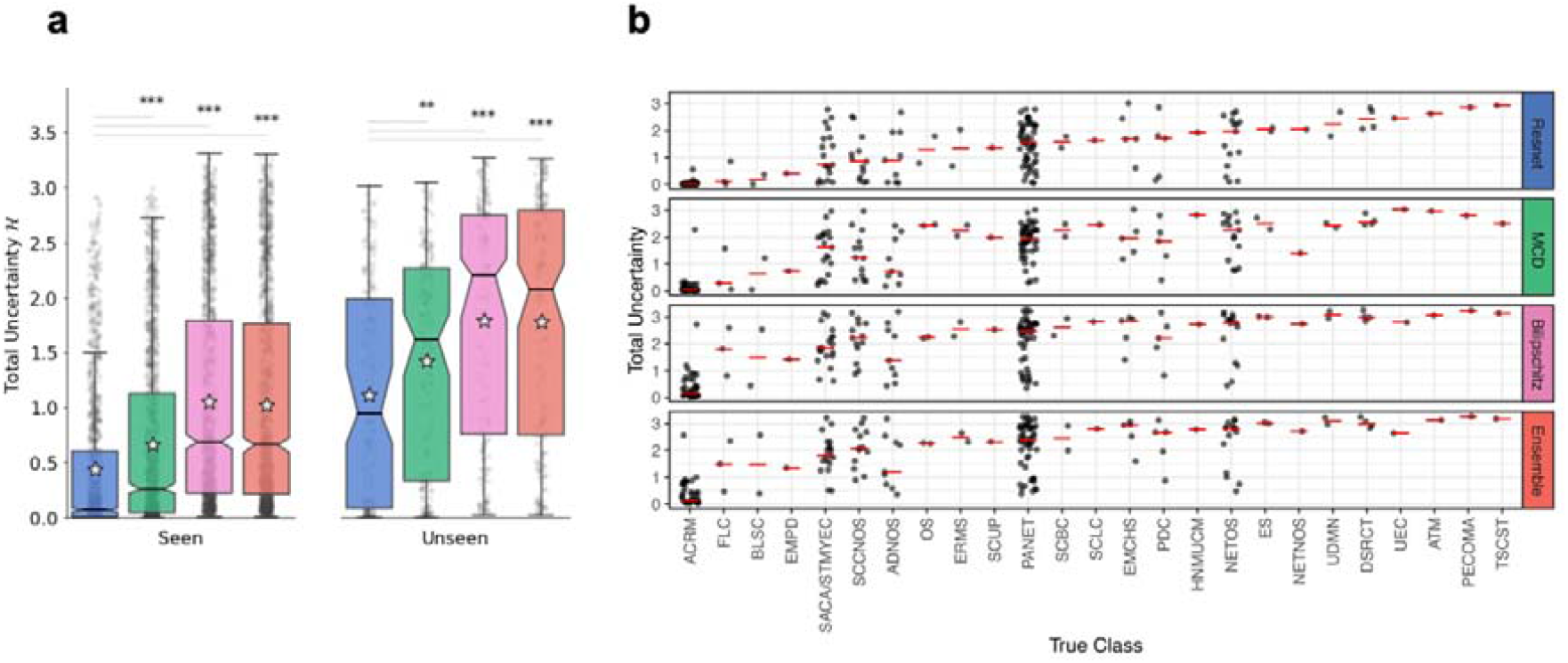
Total uncertainties for out-of-distribution data with cancer types ‘seen’ and ‘unseen’ in training. **a** Box plot of each model’s predictive uncertainty (Shannon’s Entropy, *H*) on OOD data with cancer types ‘seen’ (LHS) and ‘unseen’ (RHS) during training. Statistical significance (two-sided Wilcoxon rank-sum) between baseline and each Bayesian model are marked with denoted *, **, ***, for p-value < 0.05, p-value < 0.01, and p-value < 0.001, respectively. Stars denoted mean, the horizontal centre lines denoted median, and notches – the 95% confidence interval of the median total uncertainty. **b** Total uncertainty values for the ‘unseen’ classes. The horizontal red lines denoted median total uncertainty values.

‘Unseen’ classes (i.e. cancer types) with low levels of uncertainty (averaged within the class) corresponded to ‘seen’ classes that either (biologically) related to the predicted primary cancer type, or were from a similar tissue or cell of origin. For example, all acral melanoma (ACRM) samples (n=40), a subtype of melanoma that occurs on soles, palms and nail beds, were predicted to be cutaneous melanoma (MEL) by all four models (Supplementary Fig. 6-9) with the smallest median total uncertainty for all four models (Fig. 3b). All three fibrolamellar carcinoma (FLC) samples, a rare type of liver cancer, were predicted to be hepatocellular carcinomas (HCC), although the median uncertainty was much higher for Bilishpitz and Ensemble models compared to Resnet and MCD (1.8, 1.5, 0.1 and 0.29 Shannon’s Entropy ***H***, respectively). Two bladder squamous cell carcinomas (BLSC) showed different examples of class-level exceptions with one sample predicted as a bladder adenocarcinoma (BLCA), with the same primary tissue site as BLSC, or a lung squamous carcinoma (LUSC), with similar cell of origin. For the ‘unseen’ class pancreatic neuroendocrine tumours (PANET) we saw a wide spread of uncertainty values (Fig. 3b). Interestingly, only PANET samples that were predicted as another subtype of pancreatic cancer, pancreatic adenocarcinomas (PAAD), had low prediction uncertainty across all models compared to other incorrectly predicted PANET samples (Supplementary Fig. 10). Overall, since most of the incorrect predictions with low uncertainties had a reasonable biological explanation for the prediction, we concluded that we did not find evidence of catastrophic silent failure in this case study.

### Robustness to shift-induced overconfidence is integral for production inference

To evaluate the robustness of the models’ accuracy, as well as the uncertainty’s correlation with the error-rate (abbreviated “uncertainty’s error-rate correlation”) we used the F1-Retention Area Under the Curve (F1-AUC) [47]. Evaluation was carried out on ‘seen’ and ‘unseen’ OOD data (i.e., ‘production data’). All models yielded similar results, with only a 0.45% percent decrease between the highest and lowest ranking models (F1-AUC of 93.67% for Bilipschitz and 93.25% for MCD, respectively; Fig. 4a). The performance difference between all models was marginal as F1-AUC doesn’t capture the lost calibration caused by the distributional shift when transitioning from IID to (‘seen’ and ‘unseen’) OOD. In other words, the F1-AUC metric did not detect effects caused by the shift-induced overconfidence. This was evident from the following observations: i) inter-model accuracies were similar within IID, as well as OOD data (Fig. 2a); ii) calibration errors (i.e. overconfidence) were not different for IID (p-value > 0.05), but different for OOD (p-value < 0.01; Fig. 2d); and iii) F1-AUC scores were similar for all models, which implies ‘uncertainty’s error-rate correlation’ must have been similar (since F1-AUC encapsulates accuracy and ‘uncertainty’s error-rate correlation’ [47]). While F1-AUC encapsulated accuracy and ‘uncertainty’s error-rate correlation’, which are important components of robustness when deploying DL in production, F1-AUC does not encapsulate robustness to shift-induced overconfidence.

**Figure 4.**
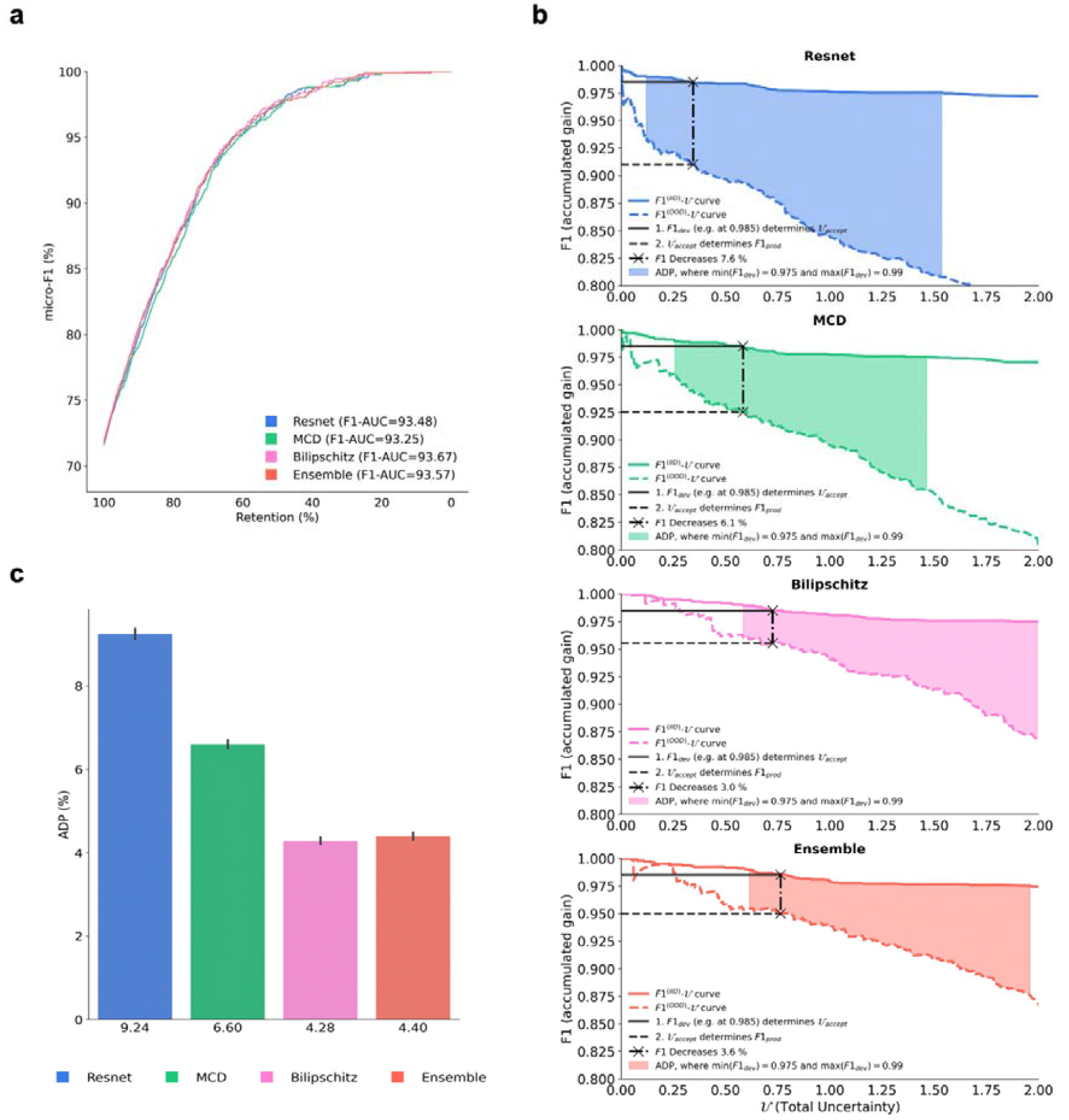
Evaluation of model generalisability from development to production. **a** F1-Retention Curves and corresponding F1-AUC scores. The F1-Retention curve of the (baseline) Resnet model and three approximate Bayesian models (MCD, Bilipschitz, Ensemble). As the retention fraction decreases, more of the most uncertain predictions are replaced with the ground truth. Thus, steeper curves require stronger correlation between uncertainty and the error-rate. The F1-Retention Area Under the Curve (F1-AUC) for each model are detailed in the legend. The F1-AUC is a function of both predictive performance (micro-F1), and the uncertainty error-rate correlation. **b** Development and Production F1-Uncertainty curves for each model. Illustrates the development F1^(IID)^-Uncertainty curves (continuous lines), as well as the production F1^(OOD)^-Uncertainty curves (dashed lines). Black lines illustrate the F1 decrease from a single development F1 score with F1_dev_=98.5 % for all models. The Area Between the Development and Production Curve (ADP) is shown as the coloured region. **c** Area Between the Development and Production Curves (ADP) bar plot with bootstrapped confidence intervals. ADP is the averaged F1 decrease calculated between F1_dev_= 97.5 % and F1_dev_= 99.0 % at intervals of 0.001 %. Steps for calculating the ADP are detailed in the Methods.

To overcome the limitation of the F1-AUC metric’s insensitivity to shift-induced overconfidence, we developed a new (prototypical) metric called the Area between the Development and Production curve (ADP), which depends on both IID (i.e. ‘development’) data, as well as the (‘seen’ and ‘unseen’) OOD (i.e. ‘production’) data. The ADP may be interpreted as “the expected decrease in accuracy when transitioning from development to production if uncertainty thresholding is utilised to boost reliability”. Furthermore, the ADP complements F1-AUC in the context of deploying models from training/development data (IID) to production test data (OOD). The ADP was calculated by averaging the set of decreases in F1, from development (IID) to production (OOD) datasets, at multiple different uncertainty thresholds (a single F1-decrease is demonstrated in Fig. 4b; refer to the Methods for details).

The ADP metric detected effects from shift-induced overconfidence, with an inter-model percent decrease that was two orders of magnitude larger than F1-AUC (Fig. 4c). The percent decrease between the top and bottom ranking models was 53.68%. The top-ranking model was Bilipschitz with an ADP of 4.28%, and the bottom ranking model was Resnet with ADP of 9.24% (Fig. 4c). This highlights that ADP may be relevant when evaluating the performance of models that are deployed in production by encapsulating shift-induced overconfidence, which is inevitable in an oncological setting.

## Discussion

A major barrier to using DL in clinical practice is the shift-induced overconfidence encountered when deploying a DL model from development to production. Reducing and accounting for shift-induced overconfidence with appropriate models and relevant metrics should make the models more transparent and trustworthy for translation into practice. Our work herein shows that marked progress can be made with simple Bayesian DL models deployed in conjunction with uncertainty thresholding. However, the performance of models deployed in production can be difficult to evaluate without a suitable metric, therefore we developed ADP to directly measure shift-induced overconfidence.

Three Bayesian models with canonical extensions, namely MCD, Bilipschitz, Ensemble, were chosen to test whether simple modifications applicable to any DL architecture can improve performance in production. The Bayesian models were selected according to the following three criteria: (1) simplicity, for wider accessibility; (2) ubiquity, to ensure models were accepted and tested methods; and (3) already demonstrated as robust to shift-induced overconfidence [22], [48], [49]. Our prior expectations were that each canonical extension would further improve generalisation of both accuracy and uncertainty quality, albeit at the cost of increased complexity. Those expectations were mostly in line with our benchmarking results, since the most complex model (Ensemble) went from worst-performing in IID to best-performing model in OOD in terms of accuracy. Furthermore, while inspection into overconfidence presented no significant inter-model differences within IID data, the OOD overconfidence differences were significant, whereby added complexity corresponded to less shift-induced overconfidence. Using the ADP statistic, improvements in robustness to shift-induced overconfidence were shown to have a large impact on the accuracy in production when rejecting unreliable predictions above an acceptable uncertainty threshold. Hence, any DL architecture’s accuracy in production can be substantially improved with simple and scalable approximate Bayesian modifications. This phenomenon is sometimes referred to as “turning the Bayesian crank” [50].

We restricted our uncertainty statistics to predictive (or total) uncertainties, since it was not possible to estimate the sub-divisions of uncertainty with the baseline Resnet model. In future work, a richer picture may be understood by focusing only on distribution-wise models to inspect the two sub-divisions of the predictive uncertainty: epistemic (model) uncertainty and aleatoric (inherent) uncertainty. Epistemic uncertainty is dependent on the model specification and may be reduced with more data or informative priors. Aleatoric uncertainty is dependent on data’s inherent noise and can be reduced with more data features that explain variance caused by confounding variables (e.g., patient age, cancer stage, batch effect). Epistemic and aleatoric uncertainties present the potential for further insights, including whether a data point’s predictive uncertainty will reduce with either more examples or by an altered model design (epistemic uncertainty), or more features (aleatoric uncertainty) [51]–[54].

This study addressed distributional shift effects on uncertainties with parametric models, which assume parameters are sufficient to represent all training data. Non-parametric models relax that assumption, which is arguably crucial to detect when data are outside the domain of training data (‘out-of-domain’) and for avoiding extreme overconfidence, i.e. ‘silent catastrophic failure’. In future work, non-parametric models, for example Gaussian Processes, capable of measuring uncertainties about ‘out-of-domain’ data, should also be explored [44]–[46], [55].

Our work suggests that considerations of robustness to distributional shifts must encapsulate uncertainty and prediction to improve performance in production. While this study focused on the quality of uncertainty, it is important to note that other DL components are worth consideration too. These include model architecture (i.e. inductive bias), which can be tailored to ignore redundant data-specific aspects of a problem via invariant or equivariant model representations [56], data-augmentation strategies [57], and/or structural causal models [58]–[60]. Such tailored models can further improve data efficiency [56], robustness to distributional shifts [27], and are central to an appropriate model specification that challenges DL deployment [61]. The importance of tailored inductive biases is supported by the prolific advances in fields beyond clinical diagnostics in computer vision (e.g. CNN’s translational equivariance [56]), and biology (e.g. how Alpha Fold 2 [62] solved the Critical Assessment of protein Structure Prediction (CASP; [63]). These studies show that a wide array of DL components can improve generalisation and thus DL performance in production. Our study argues uncertainty calibration as an important element in that array; hence, improving the quality of uncertainty can lead to improved DL reliability, in production.

In practice, we hope the community considers utilising uncertainty thresholding as a proactive method to improve accuracy and safety of DL applications, deployed in the clinic. This may involve (iterative) consultation between ML engineer and medical professionals to agree on a ‘minimally acceptable accuracy’ for production (deem this *min*(*F*1_*dev*_). The ML engineer may then use development data to train an approximate Bayesian DL model and produce Development F1-Uncertainty curves (with validation data). The engineer then, with another independent dataset, can proceed to develop an ADP estimate (as described in the methods) to help communicate (in context of available dataset differences) what the expected accuracy decrease may be when the model is deployed to production, which helps manage expectations and facilitate trust. Importantly, with the (prototypical) ADP, the team may better judge which uncertainty quantification techniques are most effective for boosting accuracy under the ‘uncertainty thresholding’ risk-management regime. This procedure, as well as the ADP statistic, is of course prototypical and only suggestive. We leave improvement, and clarification of this for future work.

In summary, we undertook this study to highlight the difficulties of DL deployment in practice, using clinical oncology as an example. We highlighted approaches for quantifying and improving robustness to shift-induced overconfidence with simple and accessible DL methods. We justified our approach with mathematical and empirical evidence, biological interpretation, and a new metric, the ADP, that is complementary to F1-AUC by encapsulating shift-induced overconfidence – a crucial aspect that needs to be considered when deploying DL in real-world production. Moreover, the ADP is directly interpretable as proxy to expected accuracy loss when deploying DL models from development to production. Although we have addressed the shift-induced overconfidence by utilising first-line solutions, work remains to bridge DL from theory to practice. We must account for data distributions, evaluation metrics, and modelling assumptions as all equally important and necessary considerations to see safe translation of DL into clinical practice.

## Methods

### Prediction task and datasets

The task was to predict a patient’s primary cancer type, which we cast under the supervised learning framework by learning the map **{x → *y*}**, with ***y*** denoting the primary cancer category, and **x ∈ *R***^***D***^denoting a patient’s sampled bulk gene expression signature.

Three independent datasets were used: our own independent Internal Custom Dataset, ICD [34]–[42], TCGA [31], and Met500 [33]. All datasets were preprocessed and partitioned into groups (i.e. strata) that uniquely proxied different distribution shifts. Proxies of approximately unique shifts were assumed to be governed by their respective intervention (i.e. unique shift), as deemed by values of four presumed hidden variables influencing the modelled map **{x → *y*}**. Those variables were *‘Batch’* (indicating source dataset label, *e*.*g*., *‘TCGA’*), *‘State-of-Metastases’* (valued *‘Primary’*, or *‘Metastatic’*), and *‘Seen’* (indicating whether a target value *y* was seen during training) (Supplementary Table 1). Training and validation data comprised of the Strata ID

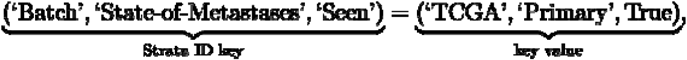

since we believed it to be approximately *independent and identically distributed* (IID) data. All other strata were assumed *out-of-distribution* (OOD) due to distribution shifts caused by confounding variables. As a result, the training and validation data were IID, while the test data were OOD.

### Benchmarked models

Four models were benchmarked in this study – the baseline pointwise Resnet, MCD, Bilipschitz, and Ensemble. All models shared identical model architecture and hyperparameter settings (including early stopping), respectively controlling the inductive bias and accuracy from confounding overconfidence.

### Baseline Resnet model

Resnet architecture had four hidden layers, each with 1024-neurons, Mish activations [64], batch normalisation [65], and standard residual connections from the first hidden layer up to the final hidden ‘logit-space’ layer, which was then normalised using the SoftMax function to yield probability vector **p(x) = ∈ [0**,**1]**^***K***^, where the prediction’s class index,

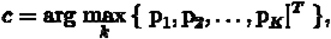

indicates the primary cancer site’s label ***y* ← *c***. Specifically, a batch **X ∈ ℝ**^***B×D***^ with ***B*** individual samples is first transformed by the input layer **U**^**(0)**^ **= *g*(⟨X**,**W**^**(0)**^**⟩ + b**^**(0)**^, with affine transform parameters {**W**^**(0)**^,**b**^**(0)**^**}**, non-linear activations ***g***, and output representation **U**^**(0)**^. Hidden layers have residual connections **U**^**(*l*)**^ **= *g*(⟨U**^**(*l* − 1)**^,**W**^**(*l*)**^**⟩ + b**^**(*I*)**^**) + U**^**(*l* − 1)**^ where ***l* ∈ 1**,**2, …, *L*** denotes the hidden layer index (***L* = 3** in this case). The final output layer is a pointwise (mean estimate) function in logit-space **f(X) = *g*(⟨U**^**(*L*)**^,**W**^**(*µ*)**^**⟩ + b**^**(*µ*)**^), where **{W**^**(*µ*)**^, **b**^**(*µ*)**^**}** are the final output (affine) transformation parameters. Finally, softmax normalisation yields a K-vector **p(X) = Softmax(f(X))**. All other hyperparameter settings are defined in Supplementary Table 2. This baseline Resnet model architecture was inherited by all other models in this study to control inductive biases.

### Approximate Bayesian inference

Bayesian inference may yield a predictive distribution about sample **x*, *p*(p**|**x\***,**𝒟)**, from the likelihood of an assumed parametric model ***p*(p**|**x*, θ)**, an (approximate) parametric posterior ***q*(θ**|**𝒟)**, and potentially Monte Carlo Integration (MCI) technique, also referred to as Bayesian model averaging:

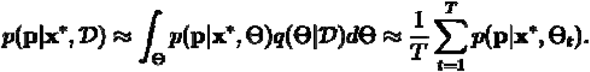

Most neural networks are parametric models, which assume **θ** can perfectly represent **𝒟**. As a result, the model likelihood ***p*(p**|**x\***,**𝒟**,**θ)** is often replaced with ***p*(p**|**x*, θ)**.The main differentiating factor among all Bayesian deep learning inference methods lies in how the parametric posterior ***q*(θ**|**𝒟)** is approximated.

### Resnet extended with Monte Carlo Dropout

The MCD model approximates the parametric posterior ***q*(θ**|**𝒟)** by keeping dropout activated during inference [43]. Dropout randomly ‘switches off’ a subset of neurons to zero-vectors at each iteration. Hence, a collection of dropout configurations 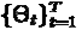 are samples from the (approximate) posterior ***q*(θ**|**𝒟)**. For more information, refer to the Appendix of [43] where an approximate dual connection between Monte Carlo Dropout neural networks and Deep Gaussian processes is established.

The MCD also extends the Resnet model architecture by including an additional output layer to estimate a data-dependent variance function 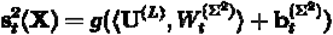 in addition to the (now stochastic) mean function 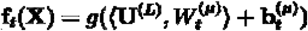. Both final output layers had a shared input **U**^**(*L*)**^, but unique parameters 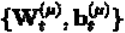 and 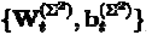. Together, the stochastic mean **f**_***i***_**(X)** and variance 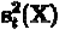 specify a Gaussian distribution in the logit-space, which was then sampled once 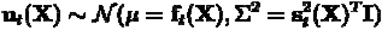 and normalised with the Softmax function **p**_***t***_**(X) = Softmax(u**_***t***_**(X)). p**_***t***_**(X)** represents a single sample from the model likelihood ***p*(p**|**x, θ)**, from which ***T*** samples are averaged for Monte Carlo integration:

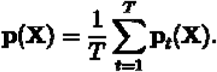

Finally, **p(X)** estimates the cancer primary site label ***y***, the predictive uncertainties **Conf(·)**, and **ℋ (·)** for each individual sample in data batch **X**.

### MCD extended with a bi-Lipschitz constraint

The BiLipschitz model shared all the properties of the MCD model with an additional bi-Lipschitz constraint:

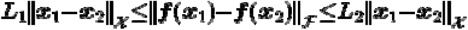

where scalars ***L***_**1**_ and ***L***_**2**_ respectively control the tightness of the lower- and upper-bound. Norm operators **{∥·∥**_**𝓍**_, **∥·∥**_**ℱ**_**}**are over the data space **𝓍** and function space **ℱ**. The effect of the bi-Lipschitz constraint is such that the changes in input data **∥x**_**1**_ **− x**_**2**_**∥**_**𝓍**_ (e.g. distribution shifts) are proportional to the changes in the output, **∥f(x**_**1**_**) − f(X**_**2**_**)∥ℱ**. These changes are within a bound determined by ***L***_**1**_ (controlling sensitivity) and ***L***_**2**_ (controlling smoothness). Interestingly, recent studies have established that bi-Lipschitz constraints are beneficial to the robustness of the neural network under distributional shifts [44], [45]. Sensitivity (i.e. ***L***_**1**_) is controlled with residual connections [66], [67], which allows **f(x)** to avoid arbitrarily small changes, especially in the presence of distributional shifts in those regions of **𝓍** with no (training data) support [44]. Sensitivity (i.e. ***L***_**2**_) is controlled with spectral normalisation on parameters **θ** [44], [68] and batch-normalisation functions [45], which allow **f(x)** to avoid arbitrarily large changes (under shifts) that induce *feature collapse* and extreme overconfidence [44]–[46].

### Deep ensemble of BiLipschitz models

The Ensemble model was a collection of eight independently trained BiLipschitz models with unique initial parameter configurations. Each Bayesian model in the Ensemble model is sampled ***T*/10(= 25)** times and then pooled to control for Monte Carlo integration between the ‘Ensemble’ and all other models.

Models in deep ensembles yield similarly performant (low-loss) solutions, but are diverse and distant in parameter- and function-space [69]. This allows the ensemble to have an (approximate) posterior ***q*(θ**|**𝒟)** with *multiple modes*, which was not the case for the Resnet, MCD, and Bilipschitz models. We believe the ensemble modelled ***q*(θ**|**𝒟)** with the highest fidelity to the true parametric posterior ***p*(θ**|**𝒟)**due to empirical evidence from other studies’ results [27], [48], [70], [71].

### Model efficacy assessment

Model efficacy was assessed using several metrics with practical relevance in mind (justification provided in the Supplementary Information – S1.2). Predictive performance, the predictive uncertainties and the *total overconfidence* were, respectively, measured with the micro-F1 score, Shannon’s Entropy***H***, and Expected Calibration Error (ECE). F1-AUC was used to evaluate the robustness of the predictive performance and the uncertainty’s error-rate correlation. The Area between Development and Production (ADP) metric was designed to complement F1-AUC by evaluating robustness to *shift-induced overconfidence*. This may be interpreted as the expected predictive loss during a model’s transition from development inference (IID) to production inference (OOD) while controlling for the uncertainty threshold.

### Quantifying predictive uncertainty

A predictive uncertainty (or total uncertainty) indicates the likelihood of an erroneous inference **p(x) = SoftMax(f(x))**, with a probability vector **p(x) = ∈ [0.1]**^***K***^, normalising operator **SoftMax(.)**, pointwise softmax function in logit-space, **f(.)**, and an gene expression vector **x ∈ ℝ**^***D***^. The ideal predictive uncertainties depend on the combination of many factors including the training data 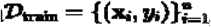, model specification (e.g. model architecture, hyperparameters, etc.), inherent noise in data, model parameters **θ**, test data inputs **x ∈ 𝒟**_**teet**_ (if modelling heteroscedastic noise), and hidden confounding variables causing distribution shifts. Consequently, there are many statistics, each explaining different phenomena, which make up the predictive uncertainty. Given that some sub-divisions of uncertainty are exclusive to distribution-wise predictive models [72], we restricted ourselves to uncertainties that are accessible to both pointwise and distribution-wise models, namely, the confidence score, **Conf(x)**, and Shannon’s Entropy ***H*(p(x))**.

A model’s confidence score w.r.t. sample **x**, is defined by the largest element from the softmax vector,

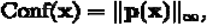

Where **p(x) = SoftMax(f(x))** and **∥p(x)∥**_**∞**_denotes the matrix-induced infinity norm of the vector **p(x)**. Confidence scores approximately quantify the probability of being correct and thus they are often used for rejecting ‘untrustworthy’ predictions (recall ‘uncertainty thresholding’ from the Introduction). Moreover, an average **Conf(x)** is comparable to the accuracy metric, which allows for evaluating the overconfidence via ECE, which we will shortly detail.

Another notion of predictive uncertainty is that of Shannon’s Entropy, i.e.,

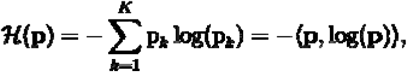

where **⟨·, ·⟩** is the dot product operator. Recall that **ℋ(p)** is maximised when **p** encodes a uniform distribution.

### Defining out-of-distribution data and the DL effects

The IID assumption on data implies true causal mechanisms (i.e. structural causal model) where the underlying data generating process is immutable across observations, and hence the samples are independently generated from the same distribution [58]. The OOD assumption, however, underpins a different setting where the underlying causal mechanisms are affected (e.g. via interventions), thus the distribution of data changes [73]. There are many different types of distributional shifts, all of which negatively affect model performance. Deep learning models can degrade under distribution shifts as the IID assumption is necessary for most optimisation strategies (Supplementary Information – S4.1). Furthermore, it is worth noting that the resulting overconfidence can be extreme, whereby arbitrary model predictions correspond with maximal confidence scores ***s***_***i***_ **→ 1** [45] (Supplementary Information – S4.2).

### Evaluation in OOD using ECE

The Expected Calibration Error was determined by binning each model’s confidence scores into ***M*** bins. The absolute difference between each bin’s accuracy and average maximum softmax score is averaged to weigh the bins proportionally with sample count. The ECE is defined as follows:

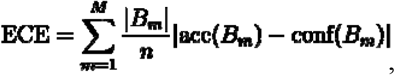

where ***B***_***m***_ is the number of predictions in bin ***m, n*** is the total number of samples, and **acc(*B***_***m***_) and **conf(*B***_***m***_) are the accuracy and confidence scores of bin ***m***, respectively.

### Evaluation in OOD using the Area under the F1-Retention Curve (F1-AUC)

Area under the F1-Retention Curve (F1-AUC) was used to evaluate model performance in OOD, as it accounts for both predictive accuracy and an uncertainty’s error-rate correlation [47]. High F1-AUC values result from high accuracy (reflected by vertical shifts in F1-Retention curves) and/or high uncertainty error-rate correlation (reflected by the gradient of the F1-Retention curves). An uncertainty’s error-rate correlation is important in the production (OOD) context as higher correlations imply more discarded erroneous predictions.

F1-AUC was quantified according to the following method.

1. Predictions were sorted by their descending order of uncertainty.
2. All predictions were iterated over in order once, while at each iteration, F1 and retention (initially 100 %) were calculated before replacing the current prediction with ground truth, hence decreasing the retention.
3. The increasing F1 scores and the corresponding decreasing retention rates determined the F1-Retention curve.
4. Approximate integration of the F1-Retention curve determined F1-AUC.

F1-Retention curves and F1-AUC metrics were quantified for all models on OOD data, including samples with classes that were not seen during training.

### Using ADP for evaluating models in OOD data relative to IID data

The Area between the Development and Production Curve (ADP) aimed to complement F1-AUC, especially in the context of deploying models from development inference (IID) to production inference (OOD). Thus, ADP was designed to capture (in OOD data, relative to IID) three aspects of a model’s robustness relating to the accuracy, uncertainty error-rate correlation, and *shift-induced overconfidence*. This is because benchmarked inter-model performance can reduce similarly in terms of robustness to accuracy and uncertainty’s error-rate correlation (as measured by F1-AUC), but *significantly differ by their uncertainty calibration* (as measured by ADP).

ADP was calculated according to the following method:

1. *Development* and *Production F1-Uncertainty curves* were produced by iteratively calculating F1 and discarding (not replacing) samples by their descending order of uncertainty.
2. A nominal F1 target range of was selected, based on the Development F1-Uncertainty curve; with **(*F*1**_***dev***_,***U***_***accept***_) denoting a point on the *Development F1-Uncertainty curve* at uncertainty threshold ***U***_***accept***_.
3. ***F*1**_***dev***_ was incremented at intervals of 1e-5 from ***F*1**_***dev***_ **= min(*F*1**_***dev***_) to ***F*1**_***dev***_ **= max(*F*1**_***dev***_), with the per cent decrease in F1, from development to production, recalculated at each step,

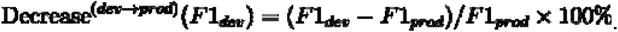
4. The set of recalculated **Decrease**^**(dev→prod)**^**(*F*1**_***dev***_) values was averaged to *approximate* the Area between the Development and Production curves (ADP).

The ADP may be interpreted as “the expected decrease in accuracy when transitioning from development to production if uncertainty thresholding is utilised to boost reliability”.

It is important to note that our method for selecting the range was not arbitrary and required two checks for each model’s Development F1-Uncertainty curve. The first check was to ensure the sample size corresponding to was sufficiently large (see Supplementary Table 3). The second check was to ensure that was large enough to satisfy production needs. *Failing to undertake these checks may result in the ADP statistic to mislead explanations about the expected loss when deploying models to production*.

ADP is practically relevant by relating to the *uncertainty thresholding* technique for improving reliability in production (recall introduction). This is because **Decrease**^**(dev→prod)**^**(*F*1**_***dev***_) first depends on a *nominated* target performance ***F*1**_***dev***_, which selects corresponding **𝒰**_***accept***_ from the Development F1-Uncertainty Curve. Predictions with uncertainties below **𝒰**_***accept***_ are accepted in production, with performance denoted by ***F*1**_***prod***_. As far as the authors are aware, no other metric monitors the three robustness components of accuracy, uncertainty’s error-rate correlation, and *shift-induced overconfidence*.

## Data availability

This project used RNA-seq data, which was previously published or available at European Genome-Phenome Archive (EGA) - EGAS00001002864. TCGA data was accessed from the National Cancer Institute Genomic Data Commons data portal (downloaded on 23rd Mar 2020), Met500 data was accessed from the University of California Santa Cruz Xena (downloaded 10th Oct 2020), and ICD data is available at EGA under study accession numbers EGAS00001000397, EGAS00001001552, EGAS00001003438, EGAS00001000154, EGAS00001001732, EGAS00001004619 and EGAS00001002864.

## Ethics approval and consent to participate

This project used RNA-seq data which was previously published or is in the process of publication. The QIMR Berghofer Human Research Ethics Committee approved use of public data (P2095).

## Code availability

Code available upon request.

## Acknowledgements

Special thanks to Yousef Rabi for their helpful discussions. We would also like to acknowledge members of the Medical Genomics and Genome Informatics teams at the QIMR Berghofer Medical Research for their technical support. This research was partially supported by the Australian Research Council through an Industrial Transformation Training Centre for Information Resilience (IC200100022). This research was also partially supported by the Cooperative Research Centres (CRC - P) Grant (CRCPFIVE000176). Nicola Waddell is supported by a National Health and Medical Research Council of Australia (NHMRC) Senior Research Fellowship (APP1139071), Olga Kondrashova is supported by a NHMRC Emerging Leader 1 Investigator Grant (APP2008631). The results published here are in whole or part based upon data generated by the TCGA Research Network: https://www.cancer.gov/tcga.

## Author contributions

M.T., O.K., & N.W. supervised the project. M.Y., H.F., R.L.J., L.T.K, O.K., & S.M pre-processed the data. S.S., O.K., H.F., M.Y., & S.M. harmonised cancer type classes. S.M., H.F., & K.S. programmed and reviewed the models. N.W., M.T., J.V.P., & F.R. provided funding for the study. J.V.P. & S.W. provided study resources. V.A. assisted with study design and results interpretation. S.M., O.K., M.T., N.W. & F.R. conceived the study with input from all other authors. S.M., O.K., M.Y., H.F., M.T., & N.W. wrote the manuscript with contributions from all other authors.

## Competing interests

M.Y., H.F., S.M., K.S. and M.T. are employed by Max Kelsen, which is a commercial company with an embedded research team. J.V.P. and N.W. are founders and shareholders of genomiQa Pty Ltd, and members of its Board. S.S., A.B., O.K., V.A., S.W, L.T.K. and R.L.J have no competing interests.

## References

[1] C. Cao et al., “Deep Learning and Its Applications in Biomedicine,” Genomics Proteomics Bioinformatics, vol. 16, no. 1, pp. 17–32, Feb. 2018, DOI: 10.1016/j.gpb.2017.07.003.

[2] K. A. Tran, O. Kondrashova, A. Bradley, E. D. Williams, J. V. Pearson, and N. Waddell, “Deep learning in cancer diagnosis, prognosis and treatment selection,” Genome Med., vol. 13, no. 1, p. 152, Sep. 2021, DOI: 10.1186/s13073-021-00968-x.

[3] M. Wang, Q. Zhang, S. Lam, J. Cai, and R. Yang, “A Review on Application of Deep Learning Algorithms in External Beam Radiotherapy Automated Treatment Planning,” Front. Oncol., vol. 10, 2020, Accessed: Jun. 06, 2022. [Online]. Available: https://www.frontiersin.org/article/10.3389/fonc.2020.580919

[4] W. Zhu, L. Xie, J. Han, and X. Guo, “The Application of Deep Learning in Cancer Prognosis Prediction,” Cancers, vol. 12, no. 3, Mar. 2020, DOI: 10.3390/cancers12030603.

[5] P. Schelb et al., “Classification of Cancer at Prostate MRI: Deep Learning versus Clinical PI-RADS Assessment,” Radiology, vol. 293, no. 3, pp. 607–617, Dec. 2019, DOI: 10.1148/radiol.2019190938.

[6] O. Ozdemir, R. Russell, and A. Berlin, “A 3D Probabilistic Deep Learning System for Detection and Diagnosis of Lung Cancer Using Low-Dose CT Scans,” IEEE Trans. Med. Imaging, vol. PP, pp. 1–1, Oct. 2019, DOI: 10.1109/TMI.2019.2947595.

[7] A. Su et al., “A deep learning model for molecular label transfer that enables cancer cell identification from histopathology images,” Npj Precis. Oncol., vol. 6, no. 1, Art. no. 1, Mar. 2022, DOI: 10.1038/s41698-022-00252-0.

[8] W. Jiao et al., “A deep learning system accurately classifies primary and metastatic cancers using passenger mutation patterns,” Nat. Commun., vol. 11, no. 1, Art. no. 1, Feb. 2020, DOI: 10.1038/s41467-019-13825-8.

[9] Z. K. Tuong et al., “Resolving the immune landscape of human prostate at a single-cell level in health and cancer,” Cell Rep., vol. 37, no. 12, p. 110132, Dec. 2021, DOI: 10.1016/j.celrep.2021.110132.

[10] M. Yap et al., “Verifying explainability of a deep learning tissue classifier trained on RNA-seq data,” Sci. Rep., vol. 11, no. 1, Art. no. 1, Jan. 2021, DOI: 10.1038/s41598-021-81773-9.

[11] A. Gayoso et al., “A Python library for probabilistic analysis of single-cell omics data,” Nat. Biotechnol., vol. 40, no. 2, Art. no. 2, Feb. 2022, DOI: 10.1038/s41587-021-01206-w.

[12] M. D. Luecken et al., “A sandbox for prediction and integration of DNA, RNA, and proteins in single cells,” presented at the Thirty-fifth Conference on Neural Information Processing Systems Datasets and Benchmarks Track (Round 2), Aug. 2021. Accessed: Jun. 06, 2022. [Online]. Available: https://openreview.net/forum?id=gN35BGa1Rt

[13] C. M. Park and J. H. Lee, “Deep Learning for Lung Cancer Nodal Staging and Real-World Clinical Practice,” Radiology, vol. 302, no. 1, pp. 212–213, Jan. 2022, DOI: 10.1148/radiol.2021211981.

[14] J. Weberpals et al., “Deep Learning-based Propensity Scores for Confounding Control in Comparative Effectiveness Research: A Large-scale, Real-world Data Study,” Epidemiol. Camb. Mass, vol. 32, no. 3, pp. 378–388, May 2021, DOI: 10.1097/EDE.0000000000001338.

[15] S. MacDonald, S. Kaiah, and M. Trzaskowski, “Interpretable AI in Healthcare: Enhancing Fairness, Safety, and Trust,” in Artificial Intelligence in Medicine: Applications, Limitations and Future Directions, Springer, Singapore, 2022, pp. 241–258.

[16] C. Rudin, “Stop explaining black box machine learning models for high stakes decisions and use interpretable models instead,” Nat. Mach. Intell., vol. 1, no. 5, Art. no. 5, May 2019, DOI: 10.1038/s42256-019-0048-x.

[17] Y. Gal, “Uncertainty in Deep Learning,” PhD, University of Cambridge, 2016.

[18] J. Gawlikowski et al., “A Survey of Uncertainty in Deep Neural Networks,” arXiv, 2107.03342, Jan. 2022. DOI: 10.48550/arXiv.2107.03342.

[19] A. Barragán-Montero et al., “Towards a safe and efficient clinical implementation of machine learning in radiation oncology by exploring model interpretability, explainability and data-model dependency,” Phys. Med. Ampmathsemicolon Biol., vol. 67, no. 11, p. 11TR01, May 2022, DOI: 10.1088/1361-6560/ac678a.

[20] A. Kristiadi, M. Hein, and P. Hennig, “Being Bayesian, Even Just a Bit, Fixes Overconfidence in ReLU Networks,” arXiv, 2002.10118, Jul. 2020. DOI: 10.48550/arXiv.2002.10118.

[21] M. Minderer et al., “Revisiting the Calibration of Modern Neural Networks,” in Advances in Neural Information Processing Systems, 2021, vol. 34, pp. 15682–15694. Accessed: Jun. 06, 2022. [Online]. Available: https://proceedings.neurips.cc/paper/2021/hash/8420d359404024567b5aefda1231af24-Abstract.html

[22] Y. Ovadia et al., “Can You Trust Your Model’s Uncertainty? Evaluating Predictive Uncertainty Under Dataset Shift,” arXiv, 1906.02530, Dec. 2019. DOI: 10.48550/arXiv.1906.02530.

[23] R. M. French, “Catastrophic forgetting in connectionist networks,” Trends Cogn. Sci., vol. 3, no. 4, pp. 128–135, Apr. 1999, DOI: 10.1016/S1364-6613(99)01294-2.

[24] S. Gupta et al., “Addressing catastrophic forgetting for medical domain expansion,” arXiv, 2103.13511, Mar. 2021. DOI: 10.48550/arXiv.2103.13511.

[25] C. Guo, G. Pleiss, Y. Sun, and K. Q. Weinberger, “On Calibration of Modern Neural Networks,” arXiv, 1706.04599, Aug. 2017. DOI: 10.48550/arXiv.1706.04599.

[26] M. E. Khan and H. Rue, “The Bayesian Learning Rule,” arXiv, 2107.04562, Mar. 2022. DOI: 10.48550/arXiv.2107.04562.

[27] A. G. Wilson and P. Izmailov, “Bayesian Deep Learning and a Probabilistic Perspective of Generalization,” Feb. 2020, DOI: 10.48550/arXiv.2002.08791.

[28] M. Divate, A. Tyagi, D. J. Richard, P. A. Prasad, H. Gowda, and S. H. Nagaraj, “Deep Learning-Based Pan-Cancer Classification Model Reveals Tissue-of-Origin Specific Gene Expression Signatures,” Cancers, vol. 14, no. 5, Art. no. 5, Jan. 2022, DOI: 10.3390/cancers14051185.

[29] J. K. Grewal et al., “Application of a Neural Network Whole Transcriptome–Based Pan-Cancer Method for Diagnosis of Primary and Metastatic Cancers,” JAMA Netw. Open, vol. 2, no. 4, p. e192597, Apr. 2019, DOI: 10.1001/jamanetworkopen.2019.2597.

[30] Y. Zhao et al., “CUP-AI-Dx: A tool for inferring cancer tissue of origin and molecular subtype using RNA gene-expression data and artificial intelligence,” EBioMedicine, vol. 61, p. 103030, Nov. 2020, DOI: 10.1016/j.ebiom.2020.103030.

[31] K. Tomczak, P. Czerwińska, and M. Wiznerowicz, “The Cancer Genome Atlas (TCGA): an immeasurable source of knowledge,” Contemp. Oncol. Poznan Pol., vol. 19, no. 1A, pp. A68–77, 2015, DOI: 10.5114/wo.2014.47136.

[32] K. A. Hoadley et al., “Cell-of-Origin Patterns Dominate the Molecular Classification of 10,000 Tumors from 33 Types of Cancer,” Cell, vol. 173, no. 2, pp. 291-304.e6, Apr. 2018, DOI: 10.1016/j.cell.2018.03.022.

[33] D. R. Robinson et al., “Integrative clinical genomics of metastatic cancer,” Nature, vol. 548, no. 7667, Art. no. 7667, Aug. 2017, DOI: 10.1038/nature23306.

[34] S. Akgül et al., “Intratumoural Heterogeneity Underlies Distinct Therapy Responses and Treatment Resistance in Glioblastoma,” Cancers, vol. 11, no. 2, Art. no. 2, Feb. 2019, DOI: 10.3390/cancers11020190.

[35] L. G. Aoude et al., “Radiomics Biomarkers Correlate with CD8 Expression and Predict Immune Signatures in Melanoma Patients,” Mol. Cancer Res., vol. 19, no. 6, pp. 950–956, Jun. 2021, DOI: 10.1158/1541-7786.MCR-20-1038.

[36] P. Bailey et al., “Genomic analyses identify molecular subtypes of pancreatic cancer,” Nature, vol. 531, no. 7592, Art. no. 7592, Mar. 2016, DOI: 10.1038/nature16965.

[37] N. K. Hayward et al., “Whole-genome landscapes of major melanoma subtypes,” Nature, vol. 545, no. 7653, Art. no. 7653, May 2017, DOI: 10.1038/nature22071.

[38] J. H. Lee et al., “Transcriptional downregulation of MHC class I and melanoma de-differentiation in resistance to PD-1 inhibition,” Nat. Commun., vol. 11, no. 1, Art. no. 1, Apr. 2020, DOI: 10.1038/s41467-020-15726-7.

[39] F. Newell et al., “Multiomic profiling of checkpoint inhibitor-treated melanoma: Identifying predictors of response and resistance, and markers of biological discordance,” Cancer Cell, vol. 40, no. 1, pp. 88-102.e7, Jan. 2022, DOI: 10.1016/j.ccell.2021.11.012.

[40] F. Newell et al., “Whole-genome sequencing of acral melanoma reveals genomic complexity and diversity,” Nat. Commun., vol. 11, no. 1, p. 5259, Oct. 2020, DOI: 10.1038/s41467-020-18988-3.

[41] A.-M. Patch et al., “Whole–genome characterization of chemoresistant ovarian cancer,” Nature, vol. 521, no. 7553, Art. no. 7553, May 2015, DOI: 10.1038/nature14410.

[42] A. Scarpa et al., “Whole-genome landscape of pancreatic neuroendocrine tumours,” Nature, vol. 543, no. 7643, Art. no. 7643, Mar. 2017, DOI: 10.1038/nature21063.

[43] Y. Gal and Z. Ghahramani, “Dropout as a Bayesian Approximation: Representing Model Uncertainty in Deep Learning,” arXiv, 1506.02142, Oct. 2016. DOI: 10.48550/arXiv.1506.02142.

[44] J. Z. Liu, Z. Lin, S. Padhy, D. Tran, T. Bedrax-Weiss, and B. Lakshminarayanan, “Simple and Principled Uncertainty Estimation with Deterministic Deep Learning via Distance Awareness,” arXiv, 2006.10108, Oct. 2020. DOI: 10.48550/arXiv.2006.10108.

[45] J. van Amersfoort, L. Smith, A. Jesson, O. Key, and Y. Gal, “On Feature Collapse and Deep Kernel Learning for Single Forward Pass Uncertainty,” arXiv, 2102.11409, Mar. 2022. DOI: 10.48550/arXiv.2102.11409.

[46] J. van Amersfoort, L. Smith, Y. W. Teh, and Y. Gal, “Uncertainty Estimation Using a Single Deep Deterministic Neural Network,” arXiv, 2003.02037, Jun. 2020. DOI: 10.48550/arXiv.2003.02037.

[47] A. Malinin et al., “Shifts: A Dataset of Real Distributional Shift Across Multiple Large-Scale Tasks,” arXiv, 2107.07455, Feb. 2022. DOI: 10.48550/arXiv.2107.07455.

[48] P. Izmailov, P. Nicholson, S. Lotfi, and A. G. Wilson, “Dangers of Bayesian Model Averaging under Covariate Shift,” in Advances in Neural Information Processing Systems, 2021, vol. 34, pp. 3309–3322. Accessed: Jun. 06, 2022. [Online]. Available: https://proceedings.neurips.cc/paper/2021/hash/1ab60b5e8bd4eac8a7537abb5936aadc-Abstract.html

[49] J. Mukhoti, P. Stenetorp, and Y. Gal, “On the Importance of Strong Baselines in Bayesian Deep Learning,” arXiv, 1811.09385, Nov. 2018. DOI: 10.48550/arXiv.1811.09385.

[50] Kevin P. Murphy, “Inference algorithms: an overview,” in Probabilistic Machine Learning: Advanced Topics (draft), MIT Press, 2022, p. 319. [Online]. Available: probml.ai

[51] M. Abdar et al., “A review of uncertainty quantification in deep learning: Techniques, applications and challenges,” Inf. Fusion, vol. 76, pp. 243–297, Dec. 2021, DOI: 10.1016/j.inffus.2021.05.008.

[52] E. Hüllermeier and W. Waegeman, “Aleatoric and epistemic uncertainty in machine learning: an introduction to concepts and methods,” Mach. Learn., vol. 110, no. 3, pp. 457–506, Mar. 2021, DOI: 10.1007/s10994-021-05946-3.

[53] A. Jesson, S. Mindermann, Y. Gal, and U. Shalit, “Quantifying Ignorance in Individual-Level Causal-Effect Estimates under Hidden Confounding,” in Proceedings of the 38th International Conference on Machine Learning, Jul. 2021, pp. 4829–4838. Accessed: Jun. 29, 2022. [Online]. Available: https://proceedings.mlr.press/v139/jesson21a.html

[54] A. S. Sambyal, N. C. Krishnan, and D. R. Bathula, “Towards Reducing Aleatoric Uncertainty for Medical Imaging Tasks.” arXiv, May 08, 2022. DOI: 10.48550/arXiv.2110.11012.

[55] S. W. Ober, C. E. Rasmussen, and M. van der Wilk, “The Promises and Pitfalls of Deep Kernel Learning,” arXiv, 2102.12108, Jul. 2021. DOI: 10.48550/arXiv.2102.12108.

[56] M. M. Bronstein, J. Bruna, T. Cohen, and P. Velickovic, “Geometric Deep Learning: Grids, Groups, Graphs, Geodesics, and Gauges.” arXiv, May 02, 2021. DOI: 10.48550/arXiv.2104.13478.

[57] C. Shorten and T. M. Khoshgoftaar, “A survey on Image Data Augmentation for Deep Learning,” J. Big Data, vol. 6, no. 1, p. 60, Jul. 2019, DOI: 10.1186/s40537-019-0197-0.

[58] J. Peters, D. Janzing, and B. Schölkopf, Elements of Causal Inference: Foundations and Learning Algorithms. Cambridge, MA, USA: MIT Press, 2017.

[59] B. Schölkopf et al., “Toward Causal Representation Learning,” Proc. IEEE, vol. 109, no. 5, pp. 612–634, May 2021, DOI: 10.1109/JPROC.2021.3058954.

[60] K. Xia, K.-Z. Lee, Y. Bengio, and E. Bareinboim, “The Causal-Neural Connection: Expressiveness, Learnability, and Inference,” in Advances in Neural Information Processing Systems, 2021, vol. 34, pp. 10823–10836. Accessed: Jun. 29, 2022. [Online]. Available: https://proceedings.neurips.cc/paper/2021/hash/5989add1703e4b0480f75e2390739f34-Abstract.html

[61] A. D’Amour et al., “Underspecification Presents Challenges for Credibility in Modern Machine Learning.” arXiv, Nov. 24, 2020. DOI: 10.48550/arXiv.2011.03395.

[62] J. Jumper et al., “Highly accurate protein structure prediction with AlphaFold,” Nature, vol. 596, no. 7873, Art. no. 7873, Aug. 2021, DOI: 10.1038/s41586-021-03819-2.

[63] A. Kryshtafovych, T. Schwede, M. Topf, K. Fidelis, and J. Moult, “Critical assessment of methods of protein structure prediction (CASP)—Round XIV,” Proteins Struct. Funct. Bioinforma., vol. 89, no. 12, pp. 1607–1617, 2021, DOI: 10.1002/prot.26237.

[64] D. Misra, “Mish: A Self Regularized Non-Monotonic Activation Function,” arXiv, 1908.08681, Aug. 2020. DOI: 10.48550/arXiv.1908.08681.

[65] S. Ioffe and C. Szegedy, “Batch Normalization: Accelerating Deep Network Training by Reducing Internal Covariate Shift,” arXiv, 1502.03167, Mar. 2015. DOI: 10.48550/arXiv.1502.03167.

[66] J. Behrmann, W. Grathwohl, R. T. Q. Chen, D. Duvenaud, and J.-H. Jacobsen, “Invertible Residual Networks,” arXiv, 1811.00995, May 2019. DOI: 10.48550/arXiv.1811.00995.

[67] K. He, X. Zhang, S. Ren, and J. Sun, “Deep Residual Learning for Image Recognition,” arXiv, 1512.03385, Dec. 2015. DOI: 10.48550/arXiv.1512.03385.

[68] F. Farnia, J. M. Zhang, and D. Tse, “Generalizable Adversarial Training via Spectral Normalization,” arXiv, 1811.07457, Nov. 2018. DOI: 10.48550/arXiv.1811.07457.

[69] S. Fort, H. Hu, and B. Lakshminarayanan, “Deep Ensembles: A Loss Landscape Perspective,” arXiv, 1912.02757, Jun. 2020. DOI: 10.48550/arXiv.1912.02757.

[70] P. Izmailov, S. Vikram, M. D. Hoffman, and A. G. Wilson, “What Are Bayesian Neural Network Posteriors Really Like?,” arXiv, 2104.14421, Apr. 2021. DOI: 10.48550/arXiv.2104.14421.

[71] F. D’ Angelo and V. Fortuin, “Repulsive Deep Ensembles are Bayesian,” in Advances in Neural Information Processing Systems, 2021, vol. 34, pp. 3451–3465. Accessed: Jun. 30, 2022. [Online]. Available: https://proceedings.neurips.cc/paper/2021/hash/1c63926ebcabda26b5cdb31b5cc91efb-Abstract.html

[72] J. Mukhoti, A. Kirsch, J. van Amersfoort, P. H. S. Torr, and Y. Gal, “Deep Deterministic Uncertainty: A Simple Baseline,” arXiv, 2102.11582, Jan. 2022. DOI: 10.48550/arXiv.2102.11582.

[73] K. Zhang, B. Schölkopf, K. Muandet, and Z. Wang, “Domain Adaptation under Target and Conditional Shift,” in Proceedings of the 30th International Conference on Machine Learning, May 2013, pp. 819–827. Accessed: Jun. 30, 2022. [Online]. Available: https://proceedings.mlr.press/v28/zhang13d.html

