## Supplementary Materials for "Generalising uncertainty improves accuracy and safety of deep learning analytics applied to oncology"

1. Max Kelsen, Brisbane, QLD, Australia
2. ARC Training Centre for Information Resilience (CIRES)
3. The University of Queensland, Brisbane, Australia
4. QIMR Berghofer Medical Research Institute, Brisbane, QLD, Australia
5. Medlab Pathology, Sydney, NSW, Australia
6. *joint senior

**Keywords:** Uncertainty, oncology, production inference, distributional shift, out-of-distribution, overconfidence, Bayesian deep learning, ML Safety

**Abstract** (149 of 150 words)

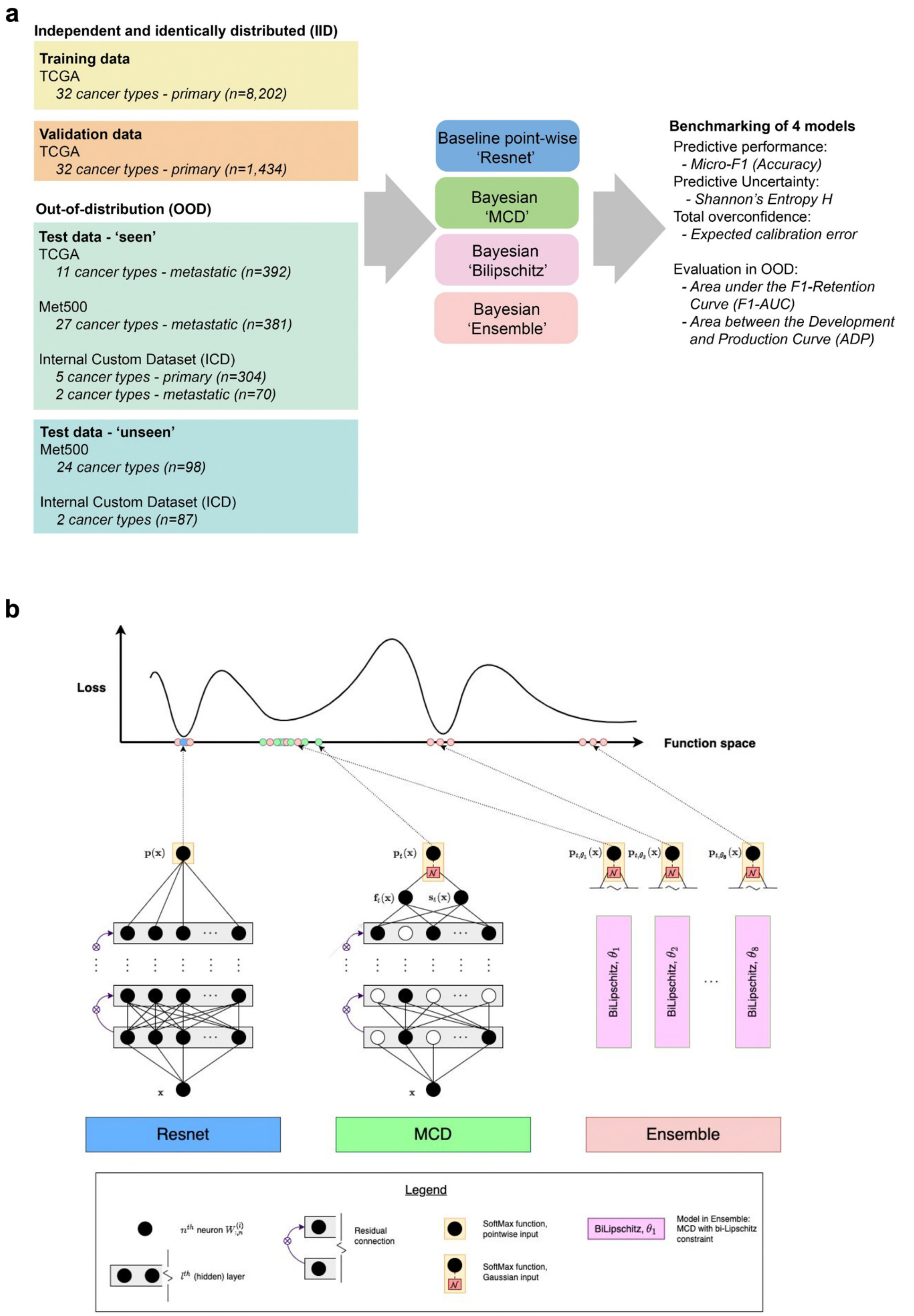


**Figure 1. Overview of the study design. a** Simplified study workflow. TCGA primary cancer types comprised the training and IID validation data. OOD test data comprised of the TCGA (metastatic cancer types), Met500 and ICD datasets, which

included primary, metastatic and 'unseen' cancer types. **b** Schematic overview of the four tested models: pointwise Resnet (Resnet), Resnet extended with Monte Carlo Dropout (MCD), MCD extended with bi-Lipschitz constraint (Bilipschitz), and an ensemble of Bilipschitz models (Ensemble). Note, Resnet represents a single point in function space (blue dot), while two Bayesian models (MCD and Bilipschitz ) represent a distribution within a single region in function space (green dots). The Ensemble represents a collection of distributions centred around different modes (red dots).

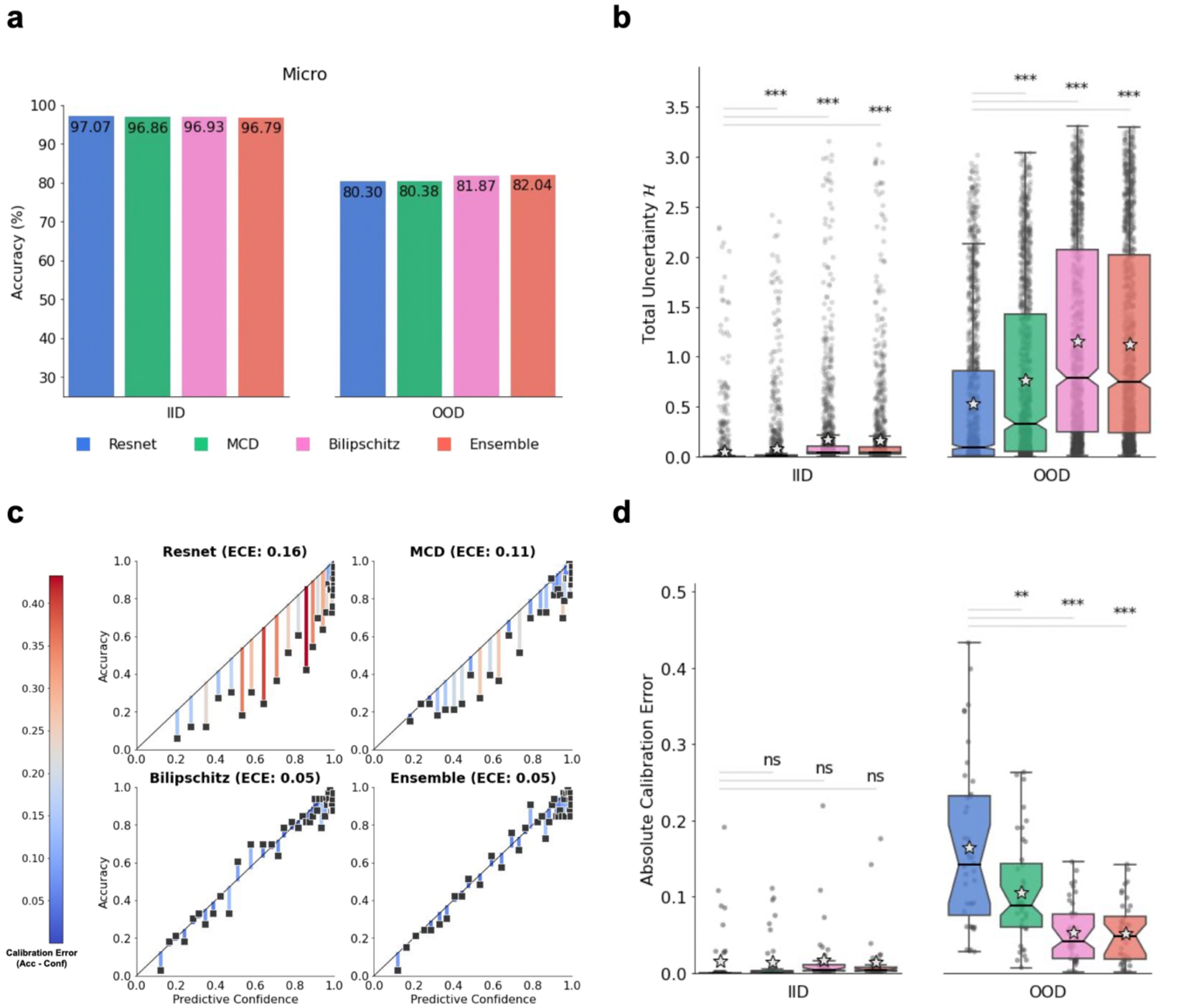


**Figure 2. Out-of-distribution overconfidence of a pointwise baseline Resnet model and three simple Bayesian models on 'seen' data. a** Micro-F1 score (i.e. Accuracy) of all models on the IID validation data (left) and on 'seen' OOD data (right). Accuracy for (IID) validation data was controlled with early stopping. **b** Box plot of each model's predictive uncertainty (Shannon's Entropy, *H*) for individual samples on IID data (left) and on 'seen' OOD data (right). Sample median is depicted by horizontal line, while the sample mean is depicted by the grey star. Statistical significance (single-sided Wilcoxon rank-sum) between baseline and each Bayesian model are marked with denoted *, **, ***, for p-value < 0.05, p-value < 0.01, and p-value < 0.001, respectively. **c** Each model's confidence vs accuracy of each ECE-bin on 'seen' OOD data. The black diagonal lines illustrate perfect calibration, i.e., no overconfidence. ECE value for each model shown in parentheses. The residuals are colour-coded by the (left) colour scale and represent the difference between confidence and accuracy for each bin. **d** Box plot of each model's absolute calibration error of individual samples on IID data (left) and 'seen' OOD data (right). Statistical significance (single-sided Wilcoxon rank-sum) between baseline and each Bayesian model are marked with denoted *, **, ***, for p-value < 0.05, p-value < 0.01, and p-value < 0.001, respectively.

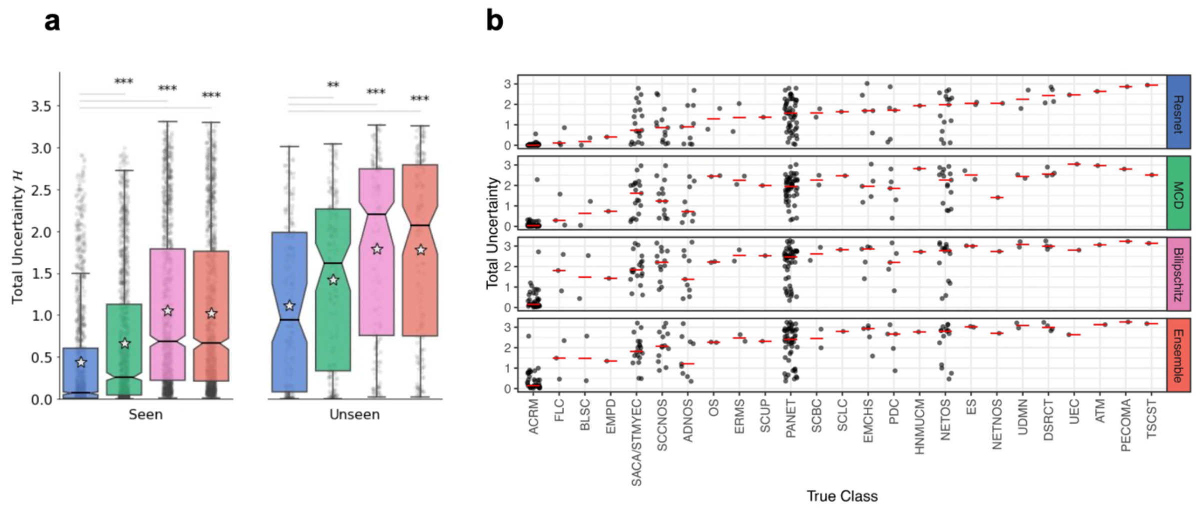


**Figure 3. Total uncertainties for out-of-distribution data with cancer types 'seen' and 'unseen' in training. a** Box plot of each model's predictive uncertainty (Shannon's Entropy, *H*) on OOD data with cancer types 'seen' (LHS) and 'unseen' (RHS) during training. Statistical significance (two-sided Wilcoxon rank-sum) between baseline and each Bayesian model are marked with denoted *, **, ***, for p-value < 0.05, p-value < 0.01, and p-value < 0.001, respectively. Stars denoted mean, the horizontal centre lines denoted median, and notches – the 95% confidence interval of the median total uncertainty. **b** Total uncertainty values for the 'unseen' classes. The horizontal red lines denoted median total uncertainty values.

#### *Robustness to shift-induced overconfidence is integral for production inference*

To evaluate the robustness of the models' accuracy, as well as the uncertainty's correlation with the error-rate (abbreviated "uncertainty's error-rate correlation") we used the F1-Retention Area Under the Curve (F1-AUC) [47]. Evaluation was carried out on 'seen' and 'unseen' OOD data (i.e., 'production data'). All models yielded similar results, with only a 0.45% percent decrease between the highest and lowest ranking models (F1-AUC of 93.67% for Bilipschitz and 93.25% for MCD, respectively; Fig. 4a). The performance difference between all models was marginal as F1-AUC doesn't capture the lost calibration caused by the distributional shift when transitioning from IID to ('seen' and 'unseen') OOD. In other words, the F1-AUC metric did not detect effects caused by the shift-induced overconfidence. This was evident from the following observations: i) inter-model accuracies were similar within IID, as well as OOD data (Fig. 2a); ii) calibration errors (i.e. overconfidence) were not different for IID (p-value > 0.05), but different for OOD (p-value < 0.01; Fig. 2d); and iii) F1-AUC scores were similar for all models, which implies 'uncertainty's error-rate correlation' must have been similar (since F1-AUC encapsulates accuracy and 'uncertainty's error-rate correlation' [47]). While F1-AUC encapsulated accuracy and 'uncertainty's error-rate correlation', which are important components of robustness when deploying DL in production, F1-AUC does not encapsulate robustness to shift-induced overconfidence.

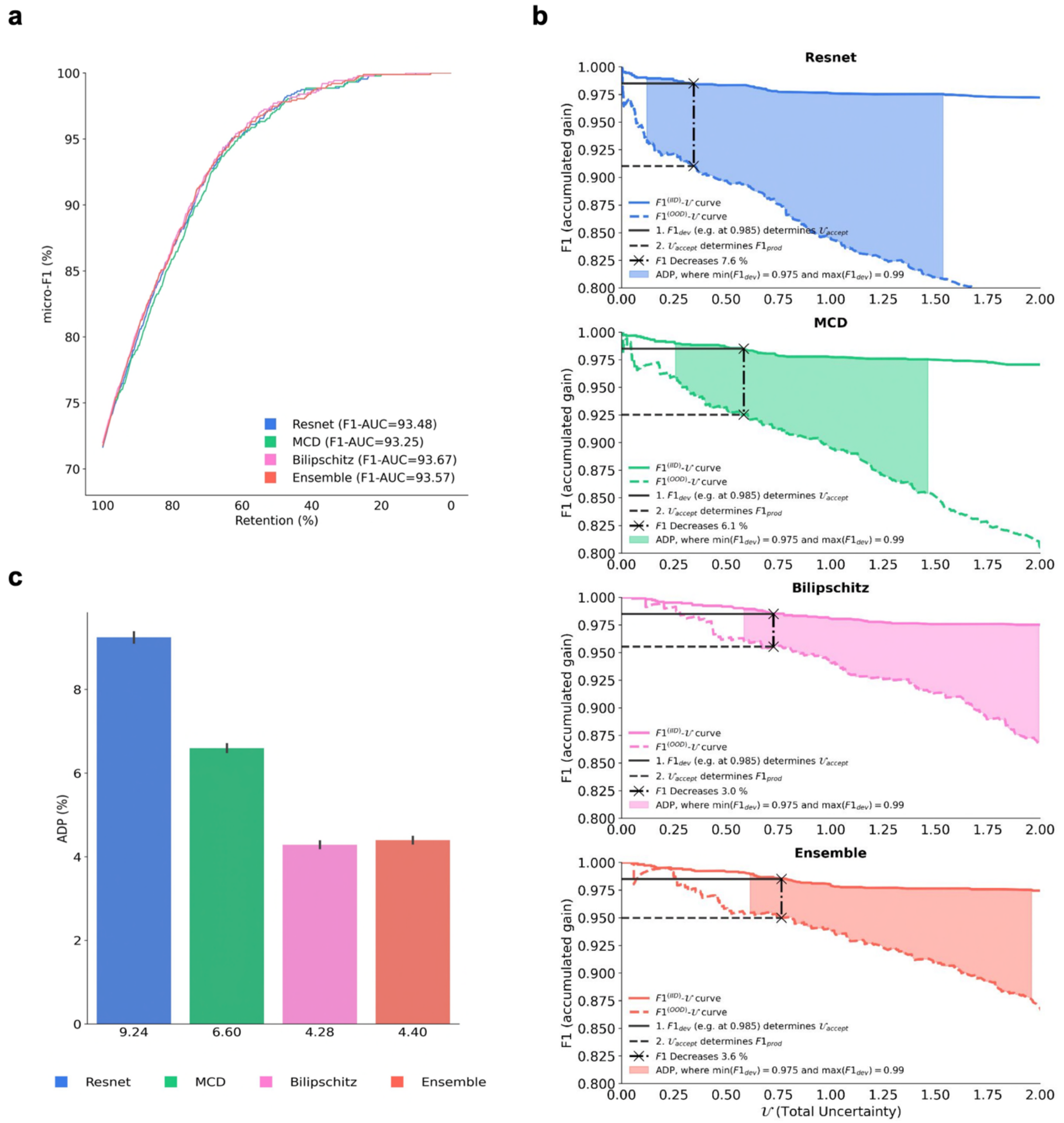


**Figure 4. Evaluation of model generalisability from development to production. a** F1-Retention Curves and corresponding F1-AUC scores. The F1-Retention curve of the (baseline) Resnet model and three approximate Bayesian models (MCD, Bilipschitz, Ensemble). As the retention fraction decreases, more of the most uncertain predictions are replaced with the ground truth. Thus, steeper curves require stronger correlation between uncertainty and the error-rate. The F1-Retention Area Under the Curve (F1-AUC) for each model are detailed in the legend. The F1-AUC is a function of both predictive performance (micro-F1), and the uncertainty error-rate correlation. **b** Development and Production F1-Uncertainty curves for each model. Illustrates the development $F1^{(IID)}$-Uncertainty curves (continuous lines), as well as the production $F1^{(OOD)}$-Uncertainty curves (dashed lines). Black lines illustrate the F1 decrease from a single development F1 score with $F1_{dev}$=98.5 % for all models. The Area Between the Development and Production Curve (ADP) is shown as the coloured region. **c** Area Between the Development and Production Curves (ADP) bar plot with bootstrapped confidence intervals. ADP is the averaged F1 decrease calculated between $F1_{dev}$= 97.5 % and $F1_{dev}$= 99.0 % at intervals of 0.001 %. Steps for calculating the ADP are detailed in the Methods.

### Methods

#### ***Prediction task and datasets***

The task was to predict a patient's primary cancer type, which we cast under the supervised learning framework by learning the map $\{\mathrm{x} \rightarrow y\}$, with $y$ denoting the primary cancer category, and $\mathrm{x} \in R^D$ denoting a patient’s sampled bulk gene expression signature.

Three independent datasets were used: our own independent Internal Custom Dataset, ICD [34]–[42], TCGA [31], and Met500 [33]. All datasets were preprocessed and partitioned into groups (i.e. strata) that uniquely proxied different distribution shifts. Proxies of approximately unique shifts were assumed to be governed by their respective intervention (i.e. unique shift), as deemed by values of four presumed hidden variables influencing the modelled map $\{\mathrm{x} \rightarrow y\}$. Those variables were *‘Batch’* (indicating source dataset label, *e.g., ‘TCGA’*), *‘State-of-Metastases’* (valued *‘Primary’*, or *‘Metastatic’*), and *‘Seen’* (indicating whether a target value *y* was seen during training) (Supplementary Table 1). Training and validation data comprised of the Strata ID

$$\underbrace{(\text{‘Batch’}, \text{‘State-of-Metastases’}, \text{‘Seen’})}_{\text{Strata ID key}} = \underbrace{(\text{‘TCGA’}, \text{‘Primary’}, \text{True})}_{\text{key value}},$$

since we believed it to be approximately *independent and identically distributed* (IID) data. All other strata were assumed *out-of-distribution* (OOD) due to distribution shifts caused by confounding variables. As a result, the training and validation data were IID, while the test data were OOD.

***Benchmarked models***

Four models were benchmarked in this study – the baseline pointwise Resnet, MCD, Bilipschitz, and Ensemble. All models shared identical model architecture and hyperparameter settings (including early stopping), respectively controlling the inductive bias and accuracy from confounding overconfidence.

***Baseline Resnet model***

Resnet architecture had four hidden layers, each with 1024-neurons, Mish activations [64], batch normalisation [65], and standard residual connections from the first hidden layer up to the final hidden 'logit-space' layer, which was then normalised using the SoftMax function to yield probability vector $\mathbf{p}(\mathbf{x}) =\in [0,1]^K$, where the prediction's class index,

$$c = \arg\max_k \{\ \mathrm{p}_1, \mathrm{p}_2, \ldots, \mathrm{p}_K]^T\ \},$$

indicates the primary cancer site's label $y \leftarrow c$. Specifically, a batch $\mathbf{X} \in \mathbb{R}^{B\times D}$ with $B$ individual samples is first transformed by the input layer $\mathbf{U}^{(0)} = g(\langle \mathbf{X}, \mathbf{W}^{(0)}\rangle + \mathbf{b}^{(0)}$, with affine transform parameters $\{\mathbf{W}^{(0)},\ \mathbf{b}^{(0)}\}$, non-linear activations $g$, and output representation $\mathbf{U}^{(0)}$. Hidden layers have residual connections $\mathbf{U}^{(l)} = g(\langle \mathbf{U}^{(l-1)}, \mathbf{W}^{(l)}\rangle + \mathbf{b}^{(l)}) + \mathbf{U}^{(l-1)}$ where $l \in 1, 2, \ldots, L$ denotes the hidden layer index ($L = 3$ in this case). The final output layer is a pointwise (mean estimate) function in logit-space $\mathbf{f}(\mathbf{X}) = g(\langle \mathbf{U}^{(L)}, \mathbf{W}^{(\mu)}\rangle + \mathbf{b}^{(\mu)})$, where $\{\mathbf{W}^{(\mu)},\ \mathbf{b}^{(\mu)}\}$ are the final output (affine) transformation parameters. Finally, softmax normalisation yields a K-vector $\mathbf{p}(\mathbf{X}) = \mathrm{Softmax}(\mathbf{f}(\mathbf{X}))$. All other hyperparameter settings are defined in Supplementary Table 2. This baseline Resnet model architecture was inherited by all other models in this study to control inductive biases.

***Approximate Bayesian inference***

Bayesian inference may yield a predictive distribution about sample $\mathbf{x}^*$, $p(\mathbf{p}|\mathbf{x}^*, \mathcal{D})$, from the likelihood of an assumed parametric model $p(\mathbf{p}|\mathbf{x}^*, \Theta)$, an (approximate) parametric posterior $q(\Theta|\mathcal{D})$, and potentially Monte Carlo Integration (MCI) technique, also referred to as Bayesian model averaging:

$$p(\mathbf{p}|\mathbf{x}^*, \mathcal{D}) \approx \int_\Theta p(\mathbf{p}|\mathbf{x}^*, \Theta) q(\Theta|\mathcal{D}) d\Theta \approx \frac{1}{T}\sum_{t=1}^{T} p(\mathbf{p}|\mathbf{x}^*, \Theta_t).$$

Most neural networks are parametric models, which assume $\Theta$ can perfectly represent $\mathcal{D}$. As a result, the model likelihood $p(\mathbf{p}|\mathbf{x}^*, \mathcal{D}, \Theta)$ is often replaced with $p(\mathbf{p}|\mathbf{x}^*, \Theta)$.The main differentiating factor among all Bayesian deep learning inference methods lies in how the parametric posterior $q(\Theta|\mathcal{D})$ is approximated.

***Resnet extended with Monte Carlo Dropout***

The MCD model approximates the parametric posterior $q(\Theta|\mathcal{D})$ by keeping dropout activated during inference [43]. Dropout randomly 'switches off' a subset of neurons to zero-vectors at each iteration. Hence, a collection of dropout configurations $\{\Theta_t\}_{t=1}^{T}$ are samples from the (approximate) posterior $q(\Theta|\mathcal{D})$. For more information, refer to the Appendix of [43] where an approximate dual connection between Monte Carlo Dropout neural networks and Deep Gaussian processes is established.

The MCD also extends the Resnet model architecture by including an additional output layer to estimate a data-dependent variance function $\mathbf{s}_t^2(\mathbf{X}) = g(\langle \mathbf{U}^{(L)}, W_t^{(\Sigma^2)} \rangle + \mathbf{b}_t^{(\Sigma^2)})$ in addition to the (now stochastic) mean function$\mathbf{f}_t(\mathbf{X}) = g(\langle \mathbf{U}^{(L)}, W_t^{(\mu)} \rangle + \mathbf{b}_t^{(\mu)})$. Both final output layers had a shared input $\mathbf{U}^{(L)}$, but unique parameters $\{\mathbf{W}_t^{(\mu)}, \mathbf{b}_t^{(\mu)}\}$ and $\{\mathbf{W}_t^{(\Sigma^2)}, \mathbf{b}_t^{(\Sigma^2)}\}$. Together, the stochastic mean $\mathbf{f}_t(\mathbf{X})$ and variance $\mathbf{s}_t^2(\mathbf{X})$ specify a Gaussian distribution in the logit-space, which was then sampled once $\mathbf{u}_t(\mathbf{X}) \sim \mathcal{N}(\mu = \mathbf{f}_t(\mathbf{X}), \Sigma^2 = \mathbf{s}_t^2(\mathbf{X})^T \mathbf{I})$ and normalised with the Softmax function $\mathbf{p}_t(\mathbf{X}) = \mathrm{Softmax}(\mathbf{u}_t(\mathbf{X}))$. $\mathbf{p}_t(\mathbf{X})$ represents a single sample from the model likelihood $p(\mathbf{p}|\mathbf{x}, \Theta)$, from which $T$ samples are averaged for Monte Carlo integration:

$$\mathbf{p}(\mathbf{X}) = \frac{1}{T} \sum_{t=1}^{T} \mathbf{p}_t(\mathbf{X}).$$

Finally, $\mathbf{p}(\mathbf{X})$ estimates the cancer primary site label $y$, the predictive uncertainties $\mathrm{Conf}(.)$, and $\mathcal{H}(.)$ for each individual sample in data batch $\mathbf{X}$.

***MCD extended with a bi-Lipschitz constraint***

The BiLipschitz model shared all the properties of the MCD model with an additional bi-Lipschitz constraint:

$$L_1 \|\boldsymbol{x}_1 - \boldsymbol{x}_2\|_{\mathcal{X}} \leq \|\boldsymbol{f}(\boldsymbol{x}_1) - \boldsymbol{f}(\boldsymbol{x}_2)\|_{\mathcal{F}} \leq L_2 \|\boldsymbol{x}_1 - \boldsymbol{x}_2\|_{\mathcal{X}}$$

where scalars $L_1$ and $L_2$ respectively control the tightness of the lower- and upper-bound. Norm operators $\{\|\cdot\|_{\mathcal{X}}, \|\cdot\|_{\mathcal{F}}\}$ are over the data space $\mathcal{X}$ and function space $\mathcal{F}$. The effect of the bi-

Lipschitz constraint is such that the changes in input data $\|\mathbf{x}_1 - \mathbf{x}_2\|_\mathcal{X}$ (e.g. distribution shifts) are proportional to the changes in the output, $\|\mathbf{f}(\mathbf{x}_1) - \mathbf{f}(\mathbf{x}_2)\|_\mathcal{F}$. These changes are within a bound determined by $L_1$ (controlling sensitivity) and $L_2$ (controlling smoothness). Interestingly, recent studies have established that bi-Lipschitz constraints are beneficial to the robustness of the neural network under distributional shifts [44], [45]. Sensitivity (i.e. $L_1$) is controlled with residual connections [66], [67], which allows $\mathbf{f}(\mathbf{x})$ to avoid arbitrarily small changes, especially in the presence of distributional shifts in those regions of $\mathcal{X}$ with no (training data) support [44]. Sensitivity (i.e. $L_2$) is controlled with spectral normalisation on parameters $\Theta$ [44], [68] and batch-normalisation functions [45], which allow $\mathbf{f}(\mathbf{x})$ to avoid arbitrarily large changes (under shifts) that induce *feature collapse* and extreme overconfidence [44]–[46].

***Deep ensemble of BiLipschitz models***

The Ensemble model was a collection of eight independently trained BiLipschitz models with unique initial parameter configurations. Each Bayesian model in the Ensemble model is sampled $T/10 (= 25)$ times and then pooled to control for Monte Carlo integration between the 'Ensemble' and all other models.

Models in deep ensembles yield similarly performant (low-loss) solutions, but are diverse and distant in parameter- and function-space [69]. This allows the ensemble to have an (approximate) posterior $q(\Theta|\mathcal{D})$ with *multiple modes*, which was not the case for the Resnet, MCD, and Bilipschitz models. We believe the ensemble modelled $q(\Theta|\mathcal{D})$ with the highest fidelity to the true parametric posterior $p(\Theta|\mathcal{D})$ due to empirical evidence from other studies' results [27], [48], [70], [71].

***Model efficacy assessment***

Model efficacy was assessed using several metrics with practical relevance in mind (justification provided in the Supplementary Information – S1.2). Predictive performance, the predictive uncertainties and the *total overconfidence* were, respectively, measured with the micro-F1 score, Shannon's Entropy $H$, and Expected Calibration Error (ECE). F1-AUC was used to evaluate the robustness of the predictive performance and the uncertainty's error-rate correlation. The Area between Development and Production (ADP) metric was designed to complement F1-AUC by evaluating robustness to *shift-induced overconfidence*. This may be interpreted as the expected predictive loss during a model's transition from development inference (IID) to production inference (OOD) while controlling for the uncertainty threshold.

#### *Quantifying predictive uncertainty*

A predictive uncertainty (or total uncertainty) indicates the likelihood of an erroneous inference $\mathbf{p}(\mathbf{x}) = \mathrm{SoftMax}(\mathbf{f}(\mathbf{x}))$, with a probability vector $\mathbf{p}(\mathbf{x}) =\in [0,1]^K$, normalising operator $\mathrm{SoftMax}(.)$, pointwise softmax function in logit-space, $\mathbf{f}(\cdot)$, and an gene expression vector $\mathbf{x} \in \mathbb{R}^D$. The ideal predictive uncertainties depend on the combination of many factors including the training data $\mathcal{D}_{\text{train}} = \{(\mathbf{x}_i, y_i)\}_{i=1}^n$, model specification (e.g. model architecture, hyperparameters, etc.), inherent noise in data, model parameters $\Theta$, test data inputs $\mathbf{x} \in \mathcal{D}_{\text{test}}$ (if modelling heteroscedastic noise), and hidden confounding variables causing distribution shifts. Consequently, there are many statistics, each explaining different phenomena, which make up the predictive uncertainty. Given that some sub-divisions of uncertainty are exclusive to distribution-wise predictive models [72], we restricted ourselves to uncertainties that are accessible to both pointwise and distribution-wise models, namely, the confidence score, $\mathrm{Conf}(\mathbf{x})$, and Shannon's Entropy $H(\mathbf{p}(\mathbf{x}))$.

A model's confidence score w.r.t. sample $\mathbf{x}$, is defined by the largest element from the softmax vector,

$$\mathrm{Conf}(\mathbf{x}) = ||\mathbf{p}(\mathbf{x})||_\infty,$$

where $\mathbf{p}(\mathbf{x}) = \mathrm{SoftMax}(\mathbf{f}(\mathbf{x}))$ and $||\mathbf{p}(\mathbf{x})||_\infty$ denotes the matrix-induced infinity norm of the vector $\mathbf{p}(\mathbf{x})$. Confidence scores approximately quantify the probability of being correct and thus they are often used for rejecting 'untrustworthy' predictions (recall 'uncertainty thresholding' from the Introduction). Moreover, an average $\mathrm{Conf}(\mathbf{x})$ is comparable to the accuracy metric, which allows for evaluating the overconfidence via ECE, which we will shortly detail.

Another notion of predictive uncertainty is that of Shannon's Entropy, i.e.,

$$\mathcal{H}(\mathbf{p}) = -\sum_{k=1}^{K} \mathrm{p}_k \log(\mathrm{p}_k) = -\langle \mathbf{p}, \log(\mathbf{p}) \rangle,$$

where $\langle \cdot, \cdot \rangle$ is the dot product operator. Recall that $\mathcal{H}(\mathbf{p})$ is maximised when $\mathbf{p}$ encodes a uniform distribution.

#### *Defining out-of-distribution data and the DL effects*

The IID assumption on data implies true causal mechanisms (i.e. structural causal model) where the underlying data generating process is immutable across observations, and hence the samples are independently generated from the same distribution [58]. The OOD assumption, however, underpins a different setting where the underlying causal mechanisms are affected (e.g. via interventions), thus the distribution of data changes [73]. There are many different types of distributional shifts, all of which negatively affect model performance. Deep learning models can degrade under distribution shifts as the IID assumption is necessary for most optimisation strategies (Supplementary Information – S4.1). Furthermore, it is worth noting that the resulting overconfidence can be extreme, whereby arbitrary model predictions correspond with maximal confidence scores $s_i \to 1$ [45] (Supplementary Information – S4.2).

***Evaluation in OOD using ECE***

The Expected Calibration Error was determined by binning each model's confidence scores into $M$ bins. The absolute difference between each bin's accuracy and average maximum softmax score is averaged to weigh the bins proportionally with sample count. The ECE is defined as follows:

$$\mathrm{ECE} = \sum_{m=1}^{M} \frac{|B_m|}{n} |\mathrm{acc}(B_m) - \mathrm{conf}(B_m)|,$$

where $B_m$ is the number of predictions in bin $m$, $n$ is the total number of samples, and $\mathrm{acc}(B_m)$ and $\mathrm{conf}(B_m)$ are the accuracy and confidence scores of bin $m$, respectively.

ADP was calculated according to the following method:

1. *Development* and *Production F1-Uncertainty curves* were produced by iteratively calculating F1 and discarding (not replacing) samples by their descending order of uncertainty.
2. A nominal F1 target range of $[\min(F1_{dev}), \max(F1_{dev})] = [0.975, 0.990]$ was selected, based on the Development F1-Uncertainty curve; with $(F1_{dev}, U_{accept})$ denoting a point on the *Development F1-Uncertainty curve* at uncertainty threshold $U_{accept}$.
3. $F1_{dev}$ was incremented at intervals of 1e-5 from $F1_{dev} = \min(F1_{dev})$ to $F1_{dev} = \max(F1_{dev})$, with the per cent decrease in F1, from development to production, recalculated at each step,

$$\text{Decrease}^{(dev \to prod)}(F1_{dev}) = (F1_{dev} - F1_{prod})/F1_{prod} \times 100\%.$$

4. The set of recalculated $\text{Decrease}^{(dev \to prod)}(F1_{dev})$ values was averaged to *approximate* the Area between the Development and Production curves (ADP).

The ADP may be interpreted as "the expected decrease in accuracy when transitioning from development to production if uncertainty thresholding is utilised to boost reliability".

It is important to note that our method for selecting the range $[\min(F1_{dev}), \max(F1_{dev})]$ was not arbitrary and required two checks for each model's Development F1-Uncertainty curve. The first check was to ensure the sample size corresponding to $\max(F1_{dev})$ was sufficiently large (see Supplementary Table 3). The second check was to ensure that $\min(F1_{dev})$ was large enough to satisfy production needs. *Failing to undertake these checks may result in the ADP statistic to mislead explanations about the expected loss when deploying models to production.*

ADP is practically relevant by relating to the *uncertainty thresholding* technique for improving reliability in production (recall introduction). This is because $\text{Decrease}^{(dev \rightarrow prod)}(F1_{dev})$ first depends on a *nominated* target performance $F1_{dev}$, which selects corresponding $\mathcal{U}_{accept}$ from the Development F1-Uncertainty Curve. Predictions with uncertainties below $\mathcal{U}_{accept}$ are accepted in production, with performance denoted by $F1_{prod}$. As far as the authors are aware, no other metric monitors the three robustness components of accuracy, uncertainty's error-rate correlation, and *shift-induced overconfidence*.
